# Gut Microbiota Regulates Exercise-induced Hormetic Modulation of Cognitive Function

**DOI:** 10.1101/2024.11.07.621818

**Authors:** Elisa Cintado, Pablo Muela, Lucia Martin-Rodriguez, Ignacio Alcaide, Patricia Tezanos, Klara Vlckova, Benjamin Valderrama, Thomaz FS Bastiaanssen, María Rodríguez-Muñoz, María L. de Ceballos, María R Aburto, John F Cryan, José Luis Trejo

## Abstract

Lifestyle factors, particularly physical exercise, significantly influence brain structure and cognitive function through a hormetic effect dependent on exercise intensity and duration. The underlying mechanisms of this profile remain largely unexplored. Recently, the gut microbiota, has emerged a potent modulator of lifestyle-induced changes on brain and behavior. Here, we demonstrate that a 40-minute protocol of moderate exercise enhances cognitive abilities related to object recognition and memory, and increases hippocampal neurogenesis in adult mice compared to sedentary controls, but these cognitive and neurogenic benefits vanish when the exercise intensity or duration is increased. Furthermore, we identified significant alpha- and beta-diversity changes and distinct bacteria composition profiles of gut microbiota associated with different exercise regimens. Specific bacterial families showed altered relative abundances depending on exercise intensity and duration, with certain families’ quantities significantly correlating with cognitive performance (*Angelakisella, Acetatifactor, Erysipelatoclostridium*, and *Coriobacteriaceae* UCG-002.). To parse causal mechanisms, fecal microbiota transplantation from exercised to sedentary mice replicated the cognitive and brain structural improvements observed in the donor animals. These findings suggest that the hormetic effects of physical exercise on cognitive function and neurogenesis are mediated by corresponding changes in the gut microbiota, indicating a novel mechanistic link between exercise, brain, and gut microbiota composition.

## Introduction

Physical exercise has been widely shown to exert widespread beneficial effects in both humans and in experimental animals ^1–5^. However, these beneficial effects obtained from moderate exercise can become detrimental when the amount, intensity, and/or duration are increased. Importantly, it is known that exercise effects follows a hormetic-like biphasic dose-response, where the biological response is increased with low doses but decreased with higher doses (inverted U-shaped curve response) ^6–9^. These beneficial effects have been reported in peripheral systems, such as the musculoskeletal or the cardiovascular systems, as well as in the nervous system. Indeed, moderate exercise is well known for its neurogenic properties in the hippocampus, which explains its pro-cognitive effects ^10–13^. Interestingly, the increase in adult hippocampal neurogenesis (AHN) promoted by exercise also appears to follow a hormetic curve ^14,15^. However, comparing different studies on exercise and its effects is challenging due to the diversity in exercise protocols, the parameters studied, and the species used in experiments. Furthermore, clear evidence of this hormetic profile has not been demonstrated so far, despite its relevance to the neurobiology of exercise. This is particularly significant because subtle changes in exercise intensity, might cause either positive or negative effects, and personalized training protocols for human subjects are lacking^16^.

Gut microbiota is one of the key components of the intestinal ecosystem and plays an essential role in health, including protection against pathogens, the configuration and maturation of immunity, regulation of metabolic intake, and absorption of nutrients and drugs ^17^. These microorganisms play also a critical role in maintaining optimal gut function and overall body health ^18,19^ through their ability to produce and release a variety of neurotransmitters, essential vitamins and amino acids, and hormones from specialized enteroendocrine cells distributed in the intestinal epithelium ^20,21^. It is largely reported that an imbalance of the gut microbiota has been associated with several disorders beyond the intestinal system. Certain pathologies are associated with dysbiosis of the gut microbiota, such as inflammatory bowel diseases ^22^, diabetes ^23^, metabolic diseases ^17^, cancer ^24^, and cardiovascular diseases ^25^. This has led to increasing investigation of therapeutic approaches, including diet, antibiotics, prebiotics, probiotics, or fecal microbiota transplantation, both to understand its function and to improve certain pathologies ^26,27^.

More recently the microbiota has emerged to play a very important role on brain and behavior. Indeed, a microbiota-gut-brain axis has been proposed and refers to the bidirectional communication between the central nervous system (CNS) and gut microbes involving a complex network of communication pathways between the endocrine system, the hypothalamic-pituitary-adrenal axis, the enteric Nervous System (ENS), and the immune system ^26,28^. Gut microbiota have been implicated in a variety of host’s CNS activities, such as cognition, adult hippocampal neurogenesis and stress response, and similarly, brain activity affects microbial composition ^29^.

In recent years, attention has focused on the relationship between lifestyle and the microbiota^30^. Given that the composition and function of the gut microbiota is highly plastic and sensitive to a wide variety of environmental factors ^31,32^, it makes the microbiota an attractive target for disease prevention and treatment, as mentioned before. Indeed, many studies have shown that different behaviors modify the microbiota, including sleep ^33^, circadian rhythms ^34^, physical activity ^35^, sedentary lifestyle ^36^, environmental enrichment ^37^ and diet ^38,39^. Recently, exercise has emerged as a powerful modulator of both the composition and metabolic activity of the gut microbiota ^40^, and the effects of chronic physical exercise have been extensively reviewed. Some of these studies have shown that regular exercise is associated with a more diverse and stable gut microbiota, which result in better gut health ^41,42^. Specific exercises like running or cycling seem to increase the abundance of specific bacterial species in humans ^43^. However, in humans, excessive exercise (in terms of intensity and/or duration) not only negatively affects the gastrointestinal system, but also alters the composition and function of the gut microbiota ^44^. In addition to increasing the abundance of beneficial bacteria, exercise has also been found to have anti-inflammatory effects on the gut by decreasing levels of pro-inflammatory cytokines ^35^. Exercise appears to also have a positive effect on gut permeability ^45,46^. Additionally, recent studies have revealed a dynamic interaction between the gut microbiota, neurodegeneration, and the role of physical activity ^35^.

In the present work we prove that varying exercise intensity protocols lead distinct changes in ANH, cognition and intestinal microbial composition. We also showed that these changes in the microbiota are sufficient to evoke said changes in neurogenesis and cognition in sedentary animals

## Results

### Study design and behavior before the physical exercise protocol

To prove whether different types of physical exercise have the same pro-cognitive effect, we implemented a longitudinal and between-groups design (SupFig. 1a and b), to evaluate the behavior of our animals both before and after the exercise treatment. The main difference between the physical exercise protocols lies in the total daily running time and the speed attained during the run, which affects the distance covered both daily (Sup.Fig. 1d) and over the entire treatment period (Sup.Fig. 1e). To clarify these differences in treatment, we represented some variables that can be seen in SupFig. 1. RUNvel (runner velocity) exercise protocol reaches the highest speeds, while the RUNtime (runner time) protocol is of longer duration. As seen in (Sup.Fig. 1d), the group that covers the longer distance per day and at the end of the treatment is the RUNtime group, followed by RUNvel, RUN (runner, moderate protocol), and finally SED (sedentary), as they do not run at all and remain in their home cage throughout the entire exercise protocol.

**Fig. 1.**
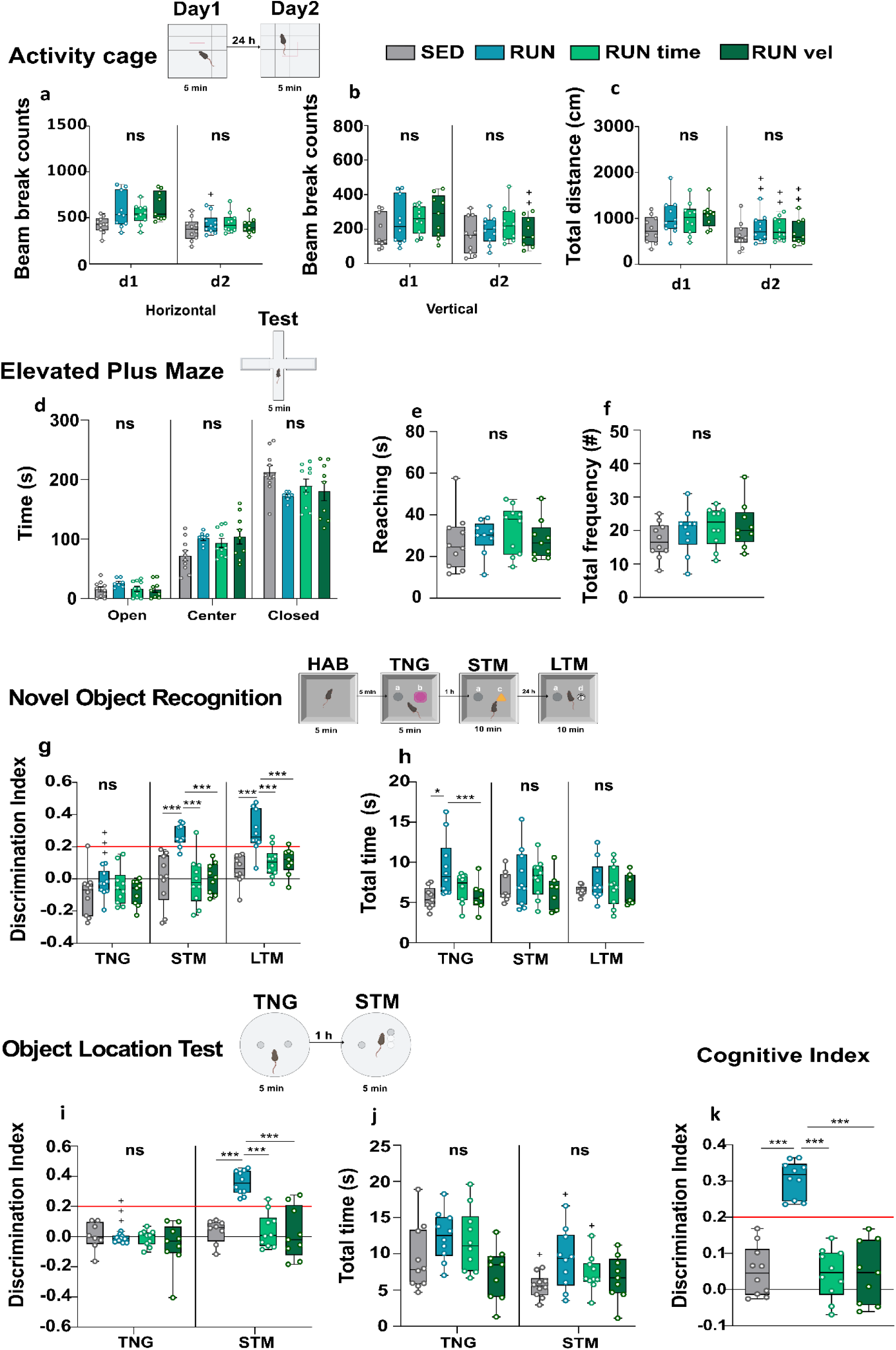
Moderate exercise induces cognitive facilitation without alterations in motor activity or anxiety-like behavior. **(a-c) Diagram of the Activity cage protocol and variables. (a) Total horizontal activity,** one-way Kruskall-wallis for each day (Day 1 (H (3, n=39) = 8.71, p-value = 0.0334); (Day 2 (H (3, n=39) = 2.01, p-value = 0.57). **(b) Total vertical activity**, Kruskall-wallis for each day (Day 1 (H (3, n=39) = 4.59, p-value = 0.204); (Day 2 (H (3, n=39) = 1.88 p-value = 0.598). **(c) Total distance traveled**, one-way ANOVA for each day (Day 1, F (1.769), p-value = 0.171), (Day 2, (F (0.587), p-value = 0.628). **(d-f) Diagram of the EPM protocol and variables. (d) Time in arms**, one-way ANOVA for the 3 phases. (Open; (F (0.446), p-value=0. 722), Center; (F (2.075), p-value=0.122), Closed; (F (2.105), p-value=0.119). **(e) Reaching,** one-way ANOVA (F (0.564), p-value=0. 643). **(f) Total frequency**, one-way ANOVA (F (1.033), p-value=0.39). **(g-h) Diagram of the NOR protocol and variables. (g) Discrimination Index in the three phases of the test.** Mixed ANOVA. Group effect (F (3,35) = 17.181, p<0.05, η2= 0.344), phase effect (F (2,70) = 34.358, p<0.05, η2= 0.387). One-way ANOVA in each phase. (TNG: F (0.74), p-value=0.532; STM: F (11.6), p-value=0.00005, η2= 0.5; LTM: F (10.9), p-value=0.00006, η2= 0.48). **(h) Total time exploration of both objects in the three phases of the test.** Group differences in each phase, one-way ANOVA and Kruskall-walis. (TNG: H (3, n=39) =11.5, p-value = 0.009, STM: F (1.11), p-value = 0.716; LTM: F (0.956), p-value = 0.716). **(i-j) Diagram of the NOL protocol and variables. (i) Discrimination Index of the columns position in both phases of the test.** Mixed ANOVA. Group effect (F (3,33) = 11.990, p<0.05, η2= 0.363), phase effect (F (1,33) = 29.995, p<0.05, η2= 0.303). One-way ANOVA in each phase. (TNG: F (0.686), p-value=0.567; STM: F (18.5), p-value=0.0000006, η2= 0.628). **(j) Total time exploration of both columns in each phase of the test.** Group differences in each phase, one-way ANOVA. (TNG: F (3.35), p-value=0.06; STM: F (2.68), p-value=0.062). **(k) Cognitive Index**. One-way ANOVA, (F (34.42), p-value=1.52e-10, η2= 0.746). Intra-group comparisons, related samples t-test. Results represent the mean ± SEM (n=10, SED group: n=10, RUN group; n=10, RUNtime group; n=9, RUNvel group). Group comparisons p-value *<0.05, **<0.01, ***<0.001. Intra-group comparisons p-value +<0.05, ++<0.01, +++<0.001. <u>Ns, not significant; TNG, training; STM, short term</u> <u>memory; LTM, long term memory.</u>

To evaluate the effect of different intensities of physical exercise on behavior, we conducted a battery of behavioral tests before and after the treatment. As expected, we confirmed that the mice started from similar baseline conditions, as indicated by the results from the activity cage (Sup. Fig. 2a-c) and the elevated plus maze (Sup. Fig. 2d-f). Additionally, we included two tests to evaluate memory: Novel Object Recognition (NOR) and Novel Object Location (NOL), to assess hippocampal-dependent memory. In the pre-treatment analysis, we confirmed that none of the experimental groups could pass these tests, as the protocols were specifically designed for control mice to not be able to pass (Sup. Fig. 2g-k). For more information, refer to the Methods section or see ^13,47^.

Therefore, if after the treatment we observe that the groups were able to pass the same test with the same level of difficulty, we could conclude that the observed cognitive facilitation was due to the physical exercise protocol.

### Moderate exercise induces cognitive facilitation without alterations in motor activity or anxiety-like behavior

Motor activity was assessed on the first day in a new environment, and on the second day in the same arena, which was now a familiar environment. As shown in (Fig. 1), we found no differences in any of the measured variables: horizontal movement (Fig. 1a), vertical movement (Fig. 1b), and distance traveled (Fig. 1c) were similar among our groups. We used an elevated plus maze test to assess anxiety-like behavior after the exercise treatment and, as shown in Figs. 1 d-f, again, we found no statistically significant differences. Indeed, we found no differences between groups, neither in the time spent in one arm or another, nor in the frequency of entries into those arms, nor in the initiative or interest in exploring the open arms (reaching).

In the NOR test, after 6 weeks of exercise, we observed that the RUN group showed a clear cognitive facilitation in this task (Fig. 1g). This group not only had higher Discrimination Index (DI) than the SED, RUNtime, and RUNvel groups in both the short-term memory (STM) phase (F (11.6), p-value< 0.00001, η2 = 0.5) and the long-term memory (LTM) phase (F (10.9), p-value< 0.00001, η2 = 0.484), but also it was the only group showing differences between the TNG (Training) and STM phases (p-value< 0.00001) and between the TNG and LTM phases (p-value< 0.000001). Therefore, they are not only significantly different from the other groups, but we also observed a significant change between the training and test phases (STM and LTM), evidencing learning.

The analysis of total exploration time in the NOR is highly relevant since it is used to calculate the DI. As shown in Fig. 1h, we do see differences in total exploration time in the training phase (TNG); TNG (H (3, n = 39) = 11.5, p-value< 0.001), but no differences in the other phases (Fig. 1 h).

In the case of the NOL test, the difficulty lies in the small separation of the objects, making it a more challenging pattern separation task. We only allowed the mice to explore both the arena and the objects for 5 minutes in each phase (Fig. 1), as in previous work ^13,47^. After the physical exercise treatment, we found that only the moderate exercise group (RUN) was able to discriminate the column position change (Fig. 1i). The RUN group had higher DIs in the STM phase (F (18.5), p-value< 0.00001), compared to the other exercised groups, while in the TNG phase, there were no differences between groups. Moreover, the DI of the RUN group in the STM phase was significantly higher than in the TNG phase (t (9) = 14.9, p-value< 0.00001). No significant differences were observed between the two phases of the test for the other groups. Regarding exploration time, as shown in Fig.1j, no differences were found between groups in either the TNG or STM phases, nor were there any intragroup differences. Therefore, the differences found in the NOL test are due to the discrimination ability itself induced by the exercise.

To summarize the differences in memory tests and to correlate with other variables we decided to calculate a cognitive index (CI), which includes the average of the three phases of both tests (STM of NOL; STM and LTM of NOR). This was intended also to filter out “false positives,”

you smooth out outlier data, or increase intra-animal consistency, or reduce variability.

as the animals would need to have performed well in all three tests to achieve a high average discrimination score. As shown in Fig. 1k, the RUN group achieved the highest CI and was significantly different from the other groups (F (34.42), p-value< 0.00000001, η² = 0.746). Therefore, with these combined behavioral results, we can affirm that the moderate exercise protocol (RUN) for 6 weeks is the only one that induces cognitive facilitation as measured in hippocampus-dependent tasks.

### Moderate exercise promotes neurogenesis and increases neural precursors and progenitors

We analyzed the volume of the hippocampal granule cell layer (GCL) and the area of the subgranular zone (SGZ). As shown in Fig. 2b and c, we did not find significant differences in either the GCL volume or the SGZ area between the different groups. Therefore, our different exercise protocols do not affect the size or volume of these two structures, which are important for memory.

**Fig. 2.**
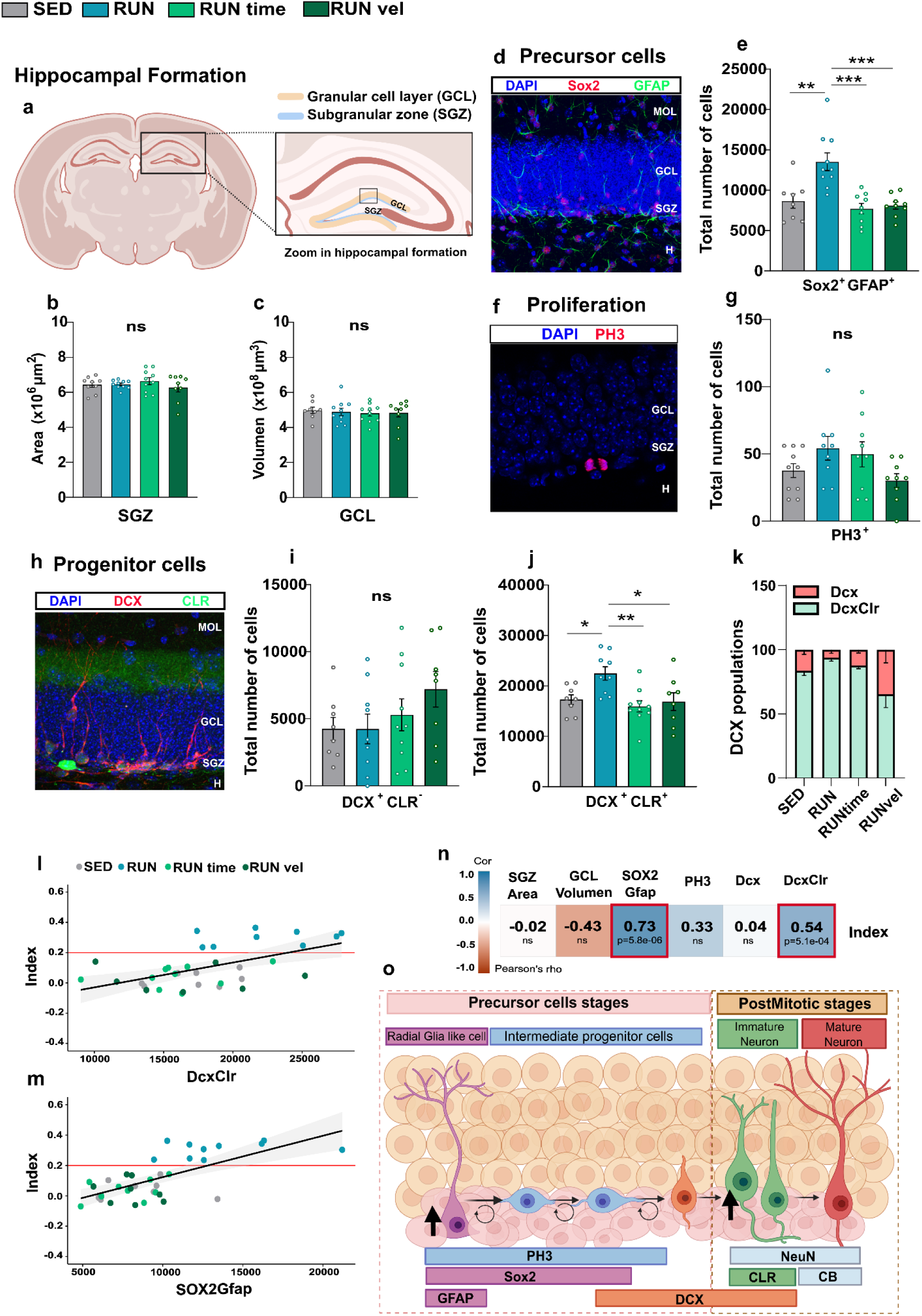
Moderate exercise promotes neurogenesis and increases neural precursors and progenitors. **(a-c)** No differences were found in the volumetric analysis of the hippocampal formation. **(a)** Schematic coronal mouse brain section indicating the structures measured within the hippocampal formation. **(b)** SGZ area, Kruskal-Wallis test, H (3, n=40) =1.19, p-value = 0.755). **(c)** GCL volumen, one-way ANOVA, F (0.25), p-value = 0.861). **(d-e)** We found some significant differences in the number of neural precursors. **(d)** Representative image of double immunofluorescence of SOX2 (red) and GFAP (green); DAPI staining is shown in blue. **(e)** Total number of SOX2^+^/GFAP^+^ cells in the SGZ, one-way ANOVA, (F (11.52), p-valor=2.78e-05, η^2^=0.519). **(f-g)** We found no differences in cell proliferation in the SGZ at the time of animal sacrifice. **(f)** Representative image of PH3+ (red) and DAPI (blue) immunofluorescence. **(g)** Total number of PH3+ cells, one-way ANOVA, (F (2.248), p-valor=0.101). **(h-k)** We found changes in certain populations of neural progenitors depending on physical exercise. **(h)** Representative image of double immunofluorescence of DCX (red) and CLR (green); DAPI staining is shown in blue. **(i)** Total number of DCX^+^/CLR^−^ cells in the GCL, one-way ANOVA, (F (1.375), p-valor=0.265). **(j)** Total number of DCX^+^/CLR^+^ cells in the GCL, one-way ANOVA with White’s adjustment, (F (6.104), p-valor=0.002, η^2^=0.376). **(k)** Proportion of DCX populations. **(l-m)** We found high correlations between precursor and progenitor populations with cognitive performance. **(l)** Correlation of the DCX^+^/CLR^+^ population with the cognitive index (r = 0.54, p-value=5.1e-4). **(m)** Correlation of the SOX2^+^/GFAP^+^ population with the cognitive index (r = 0.73, p-value = 5.8e-6). **(n)** Correlations of all NHA parameters with the cognitive index. Red boxes represent significant correlations. **(o)** Summary diagram of the cell populations studied and their histological markers. Statistical outliers were removed from the analysis. Intra-group comparisons, paired t-test or Wilcoxon test. The results represent the mean ± SEM (n=10, SED group: n=10, RUN group; n=10, RUNtime group; n=9, RUNvel group). MOL: molecular layer of the DG; SGZ: subgranular zone of the DG; H: hilus. Between-group comparisons p-value *<0.05, **<0.01, ***<0.001. Ns, not significant.

Given that the positive effects of exercise on learning and memory are mediated, among other factors, by AHN we considered its analysis indispensable in this work. To that end, we examined different markers of the neurogenic niche following the different exercise protocols. First, we analyzed whether potential changes in the neurogenic rate could be associated with variations in the number of neural precursors by performing double immunofluorescence (IF) labelling for SOX2 and GFAP. SOX2 IF labels the somas of neural precursors, while GFAP visualizes the characteristic dendritic morphology of these cells (Fig. 2d). As shown in Fig. 2e, the number of neural precursors differed significantly among the groups (F (11.52), p-value< 0.0001, η2 = 0.519), with the RUN group having a significantly higher number of neural precursors compared to the other groups.

Secondly, we analyzed the rate of proliferative cells in the DG of the hippocampus using PH3 IF, a marker of proliferation that detects cells in mitosis at the time of sacrifice (Fig. 2f-g). We found no statistical differences between groups (Fig. 2g), indicating that 6 weeks of exercise does not alter cell proliferation.

Next, we performed double IF labeling for DCX and CLR to investigate different populations of neural progenitors (Fig. 2o). This double labeling allows the identification of distinct neural progenitor populations: DCX+/CLR-cells identify type 2b neural progenitors (highly proliferative state) and type 3 progenitors (cells in a transitional state between potentially proliferative cells and post-mitotic immature neurons); DCX+/CLR+ cells correspond to differentiating immature neurons, which are post-mitotic cells. The DCX^+^/CLR^−^ population (Fig. 2i), appeared to increase with the intensity of physical exercise, although this result was not statistically significant (H (6.29) p-value=0.0981). The major change we found in neural progenitors occurred in the DCX^+^/CLR^+^ population (F (6.104), p-value=0.002), as shown in Fig. 2j. Neural progenitors increased in the RUN group compared to the other groups.

Finally, aiming to relate the analyzed parameters of the adult neurogenic niche to cognitive performance, we correlated the CI (see previous section) with all observed neurogenic markers (Fig. 2n) and found that the only significant correlations were with the precursors (Fig. 2m) and with the progenitors (Fig. 2l). Thus, high levels of cellular precursors and neural progenitors are related to better performance in hippocampus-dependent tasks.

In summary, six weeks of moderate exercise (RUN group) increased neural precursors and progenitors in the hippocampus, which correlated with cognitive improvement, whereas other types of exercise did not.

### Study of Tight Junctions and the glial cells in the Hippocampus of exercised mice

Some studies have reported changes in brain vascular tight junctions and glial cells following different exercise protocols. Therefore, we examined the presence of tight junctions in brain vessels using double IF, labeling vessels with anti-podocalyxin and tight junctions with zonula occludens 1 (ZO-1) antibodies (Fig. 3a). We found no differences in the brain area covered by vessels (Fig. 3b) across the different exercise protocols. Similarly, the area positive for ZO-1 (Fig. 3c) was comparable across groups. However, when we specifically assessed the ZO-1 positive signal within the vessels we find significant differences between groups (F (5.089), p-value = 0.009, η2= 0.432), there was a trend towards an increase in the RUN group compared to the SED and RUNtime groups, with a significant difference compared to the RUNvel group.

**Fig. 3.**
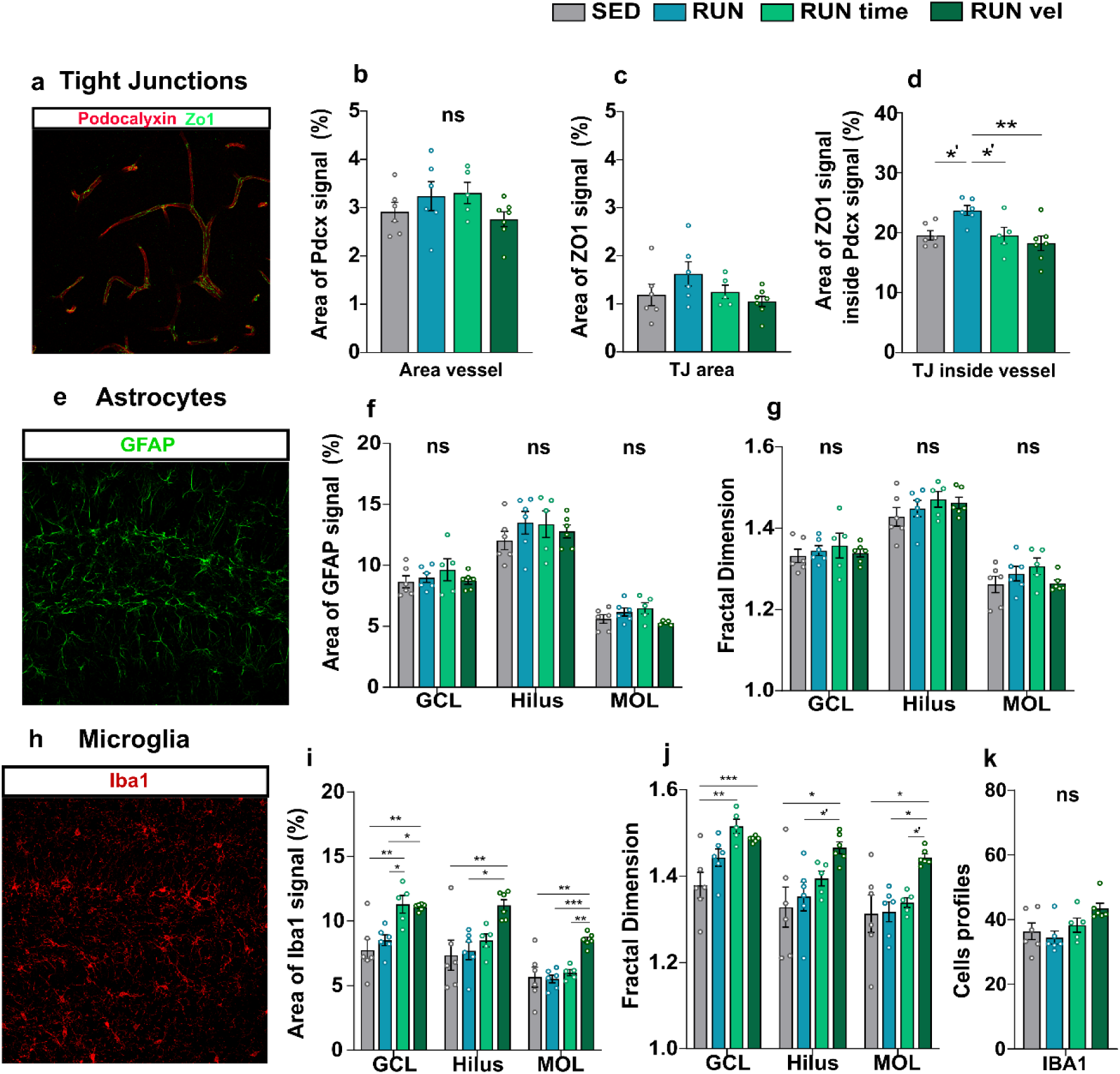
Study of Tight Junctions and Glial cells in the Hippocampus. **(a-d)** Measurement of ZO1 by Inmunofluerescence in the Hippocampus. **(a)** Representative image of IF for Pdcx (red) and ZO1 (green). **(b)** Area occupied by vessels (Pdcx). One-way ANOVA: F (1.404), p-value = 0.271. **(c)** Area occupied by ZO1. One-way ANOVA: F (1.774), p-value = 0.184. **(d)** Area occupied by ZO1 within the vessels. One-way ANOVA: F (5.089), p-value = 0.009, η2= 0.432. **(e-g)** Measurement of GFAP by IF in the Hippocampus. **(e)** Representative image of IF for GFAP (green). **(f)** Area occupied by astrocytes (GFAP). GCL; One-way ANOVA: F (0.667), p-value = 0.583. Hilus; One-way ANOVA: F (0.67), p-value=0.581. ML; One-way ANOVA: F (2.428), p-value=0.099. **(g)** Fractal dimension (GFAP). GCL; One-way ANOVA: F (0.667), p-value=0.583. Hilus; One-way ANOVA: F (0.67), p-value=0.581. ML; One-way ANOVA: F (0.896), p-value=0.462. **(h-k)** Measurement of Iba1 by IF in the Hippocampus. **(h)** Representative image of IF for Iba1 (red). **(i)** Area occupied by microglia (Iba1). GCL; One-way ANOVA: F (9.488), p-value=0.000482, η2= 0.599. Hilus; One-way ANOVA: F (5.388), p-value=0.0074, η2= 0.459. ML; One-way ANOVA with White’s correction: F (25.47), p-value=7.19e-7. **(j)** Fractal dimension (Iba1). GCL; One-way ANOVA: F (8.376), p-value=0.000942, η2= 0.569. Hilus; One-way ANOVA: F (3.788), p-value=0.027, η2= 0.374. ML; One-way ANOVA with White’s correction: F (6.129), p-value=0.0099. **(k)** Quantification of microglial cell profiles in the hippocampus. Kruskal-Wallis, H (3,24) = 6.08, p= 0.108. Results are presented as mean ± SEM (n=6, SED group; n=6, RUN group; n=6, RUNtime group; n=6, RUNvel group). Group comparisons p-value *<0.05, **<0.01, ***<0.001. Ns, not significant.

Astrocytes, labeled with GFAP, were analyzed in several hippocampal regions (Fig. 3e-g). We detected no differences in the GFAP-positive area between the groups, nor were there any differences in the fractal dimension, an index of structural complexity. Next, we stained the sections for Iba1, a marker for microglial cells, the resident macrophages in the brain. We found a significant increase in both the Iba1 signal and fractal dimension in the RUNvel group compared to the SED group, and occasionally between the RUNtime and SED groups, even though no differences in cell numbers were observed.

In summary, moderate exercise appears to strengthen the blood-brain barrier, as indicated by the increase in ZO-1 in brain vessels. While exercise does not seem to affect astrocytes, intense exercise appears to induce a mild activation of microglial cells, whereas moderate exercise does not have this effect.

### Moderate exercise is associated with increased microbiota diversity

There is accumulating evidence that the microbiota may play a role in certain behaviors and mental health in general, and it has been reported that exercise itself modifies microbial composition in mice and humans as well.

In Figures 4a-c, alpha diversity indexes are depicted: Chao1 index which estimates the number of different species in a sample; Shannon index reflects how many different taxa there are and how evenly they are distributed within a sample, and Simpson index describes even species dominance. These results revealed that the abundance of genera was different between groups (F (5.283), p-value = 0.011, η2 = 0.514), being higher in the RUN group. The moderate exercise group has a higher Chao1 index compared to RUNvel (p = 0.01) and shows a trend with the sedentary group (p = 0.059) and with the RUNtime group (p = 0.054) (Fig. 4a). This result is similar to that observed in the Shannon index, where we observed significant differences between groups (F (3.541), p-value = 0.041, η2 = 0.415), but it only becomes significant between the RUN group and the RUNvel group (p = 0.03) (Fig4b). Finally, we did not found differences between groups in the Simpson index (F (3.16), p-value = 0.056) (Fig48c). In Fig. 4d, we illustrate the differences found in beta diversity in microbial composition using Euclidean distance, presented as a Principal Component Analysis (PCA) plot (PERMANOVA: F (2.8), p-value = 0.001). The taxonomic analysis of the microbiota between groups is presented in Fig. 4e and f, where a stacked bar graph of the 20 families and the 33 most abundant genera in all groups is shown. Among the top 20 families, we observe several biologically relevant taxa, including *Bacteroidaceae, Prevotellaceae, Lachnospiraceae, and Rikenellaceae*.

**Fig.4.**
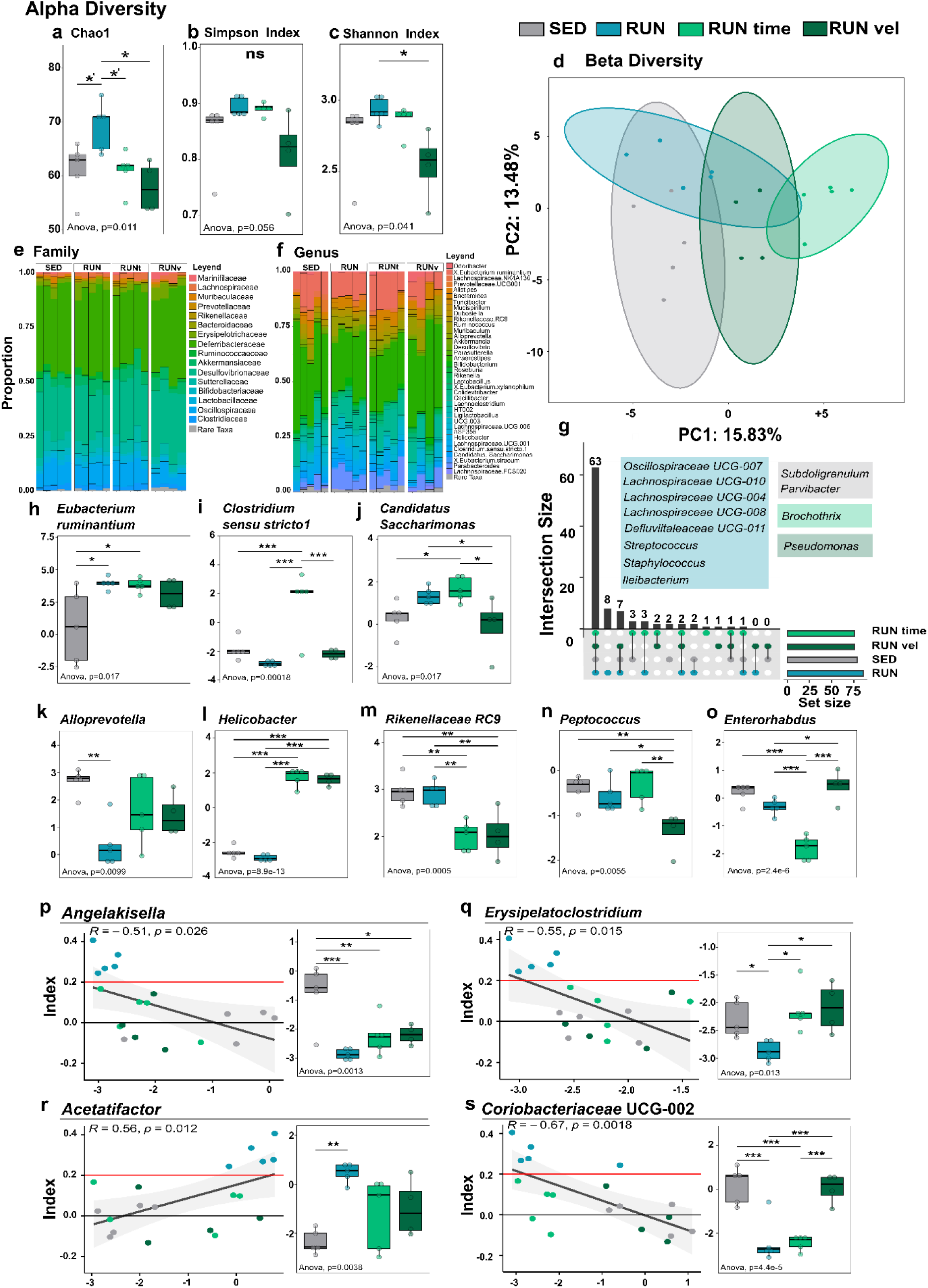
Microbial composition after physical exercise. **(a-c)** Alpha Diversity. **(a)** Chao1. One-way ANOVA: F (5.283), p-value=0.011, η2= 0.514. **(b)** Simpson Index. One-way ANOVA: F (3.16), p-value=0.056, η2= 0.387. **(c)** Shannon Index. One-way ANOVA: F (3.541), p-value=0.041, η2= 0.415. **(d)** Beta Diversity. PERMANOVA: F (2.8), p-value=0.001. **(e-f)** Taxonomy proportion by groups. Rare taxa includes all families or genera representing < 1% of the total for each sample or animal. **(e)** Proportion of Families. **(f)** Proportion of Genera. **(g)** UPSET plot of the genera present in each group out of the 96 identified genera. Intersection size represents the number of common genera among groups. **(h-o)** Differential expression of the 15 bacterial genera with fdr <=0.1. **(h)** *Eubacterium ruminantium*, One-way ANOVA: F (4.66), p-value=0.017. **(i)** *Clostridium sensu stricto1*. One-way ANOVA: F (13.08), p-value=0.00018. **(j)** *Candidatus Saccharimonas*. One-way ANOVA: F (4.67), p-value=0.016. **(k)** *Alloprevotella*. One-way ANOVA: F (5.425), p-value=0.009. **(l)** *Helicobacter*. One-way ANOVA: F (231.28), p-value=8.87e-13. **(m)** *Rikenellaceae RC9*. One-way ANOVA: F (10.78), p-value=0.0004. **(n)** *Peptococcus*. One-way ANOVA: F (6.317), p-value=0.005. **(o)** *Enterorhabdus*. One-way ANOVA: F (27.893), p-value=2.36e-6. **(p-r)** Correlation of differential expression with the Cognitive Index. **(p)** *Angelakisella*. One-way ANOVA: F (8.88), p-value=0.001. Correlation with CI, rho=-0.51, p=0.026. **(q)** *Erysipelatoclostridium*. One-way ANOVA: F (5.046), p-value=0.01. Correlation with CI, rho=-0.55, p=0.015. **(r)** *Acetatifactor*. One-way ANOVA: F (6.91), p-value=0.003. Correlation with CI, rho=0.56, p=0.12. **(s)** *Coriobacteriaceae* UCG-002. One-way ANOVA: F (16.96), p-value=4.37e-5. Correlation with CI, rho=-0.67, p=0.0018. Between-group comparisons p-value *<0.05, **<0.01, ***<0.001. Ns, not significant.

We identified 96 genera, and most of those genera are shared by all four groups (a total of 63 genera), shown in Fig. 4g. We observe that RUN group has the highest number of genera, followed by the SED group, RUNvel group, and lastly, RUNtime group. We found that there were 8 genera present only in the RUN group (*Oscillospiraceae* UCG-007, *Lachnospiraceae* UCG-004, *Lachnospiraceae* UCG-008, *Lachnospiraceae* UCG-010, *Defluvitaleaceae* UCG-011, *Streptococcus, Staphylococcus, and Ileibacterium*), 2 genera in the SED group (*Subdoligranulem, Parvibacter*), *Brochotrix* in RUNtime, and *Pseudomonas* in RUNvel. However, these genera, being so low in abundance, did not pass the filter we applied, with only 4 genera from the RUN group passing it: *Lachnospiraceae* UCG-008, *Lachnospiraceae* UCG-010, *Defluvitaleaceae* UCG-011, and *Staphylococcus*.

We then performed a differential expression analysis of bacterial genera identified in our four groups. We found approximately 15 differentially expressed genera (Fig. 4h-s) with a false discovery rate (FDR) of less than 0.1: [*Eubacterium] ruminantium group, Clostridium sensu stricto 1, Candidatus Saccharimonas, Alloprevotella, Helicobacter, Rikenellaceae* RC9 gut group, *Peptococcus, Enterorhabdus, Angelakisella, Erysipelatoclostridium, Acetatifactor, Parasutterella* (not shown), *Faecalibaculum* (not shown), *Parabacteroides* (not shown), and *Coriobacteriaceae* UCG-002.

Finally, we focused on finding correlations between cognitive performance and microbiota at the genus level, correlating the differential expression of each genus with the cognitive index to establish possible predictions of cognitive improvement. We found several candidates of interest, as shown in Figure 4 p-s, where 4 genera appear to have a high and significant correlation with the CI: *Angelakisella, Acetatifactor, Erysipelatoclostridium*, and *Coriobacteriaceae* UCG-002. In all cases, the correlations were moderate or strong (rho values of −0.51, −0.55, 0.56, and −0.63, respectively), and all were statistically significant (p < 0.05). Specifically, *Angelakisella, Erysipelatoclostridium, and Coriobacteriaceae* show negative correlations, meaning that worse cognitive performance is related to higher levels of these genera. On the other hand, *Acetatifactor* shows a positive correlation, although there are no significant differences between the RUN group and the intense exercise groups in the abundance of this genera, a trend is observed with RUNtime, while no differences are seen with RUNvel (RUN-RUNtime, p = 0.07; RUN-RUNvel, p = 0.17).

In summary, the RUN group presents greater microbial diversity, evidenced by greater alpha and beta diversity compared to the other groups. We observed that microbial composition differs not only due to exercise practice but also by the type of exercise performed. Additionally, we found a microbial profile associated with a pro-cognitive exercise protocol, specifically, the genera exclusive to this group: *Lachnospiraceae* UCG-008, *Lachnospiraceae* UCG-010, *Defluvitaleaceae* UCG-011, and *Staphylococcus*. Lastly, we identified a total of 15 genera whose expression is differentially affected by exercise practice (all genera whose differential expression has an FDR < 0.1), 4 are of particular interest as they correlate with cognitive performance.

### Fecal microbiota transplant from moderate runners induces cognitive improvement and promotes neurogenesis

To demonstrate the possible role of microbiota changes in the cognitive facilitation after moderate exercise we performed fecal microbiota transplant (FMT). The study design is depicted in Fig. 5a. To that end, 3 different groups were studied: sedentary mice; mice that did not exercise and that received a FMT from sedentary mice (SED), sedentary mice that received a FMT from mice following moderate running for 6 weeks (RUN) or from mice exercised for longer time (RUN time). After 1 week receiving an antibiotic cocktail, the FMT proceeded during week 2, and weeks 3 and 4 were devoted to behavioral testing (Fig. 5a).

**Fig. 5.**
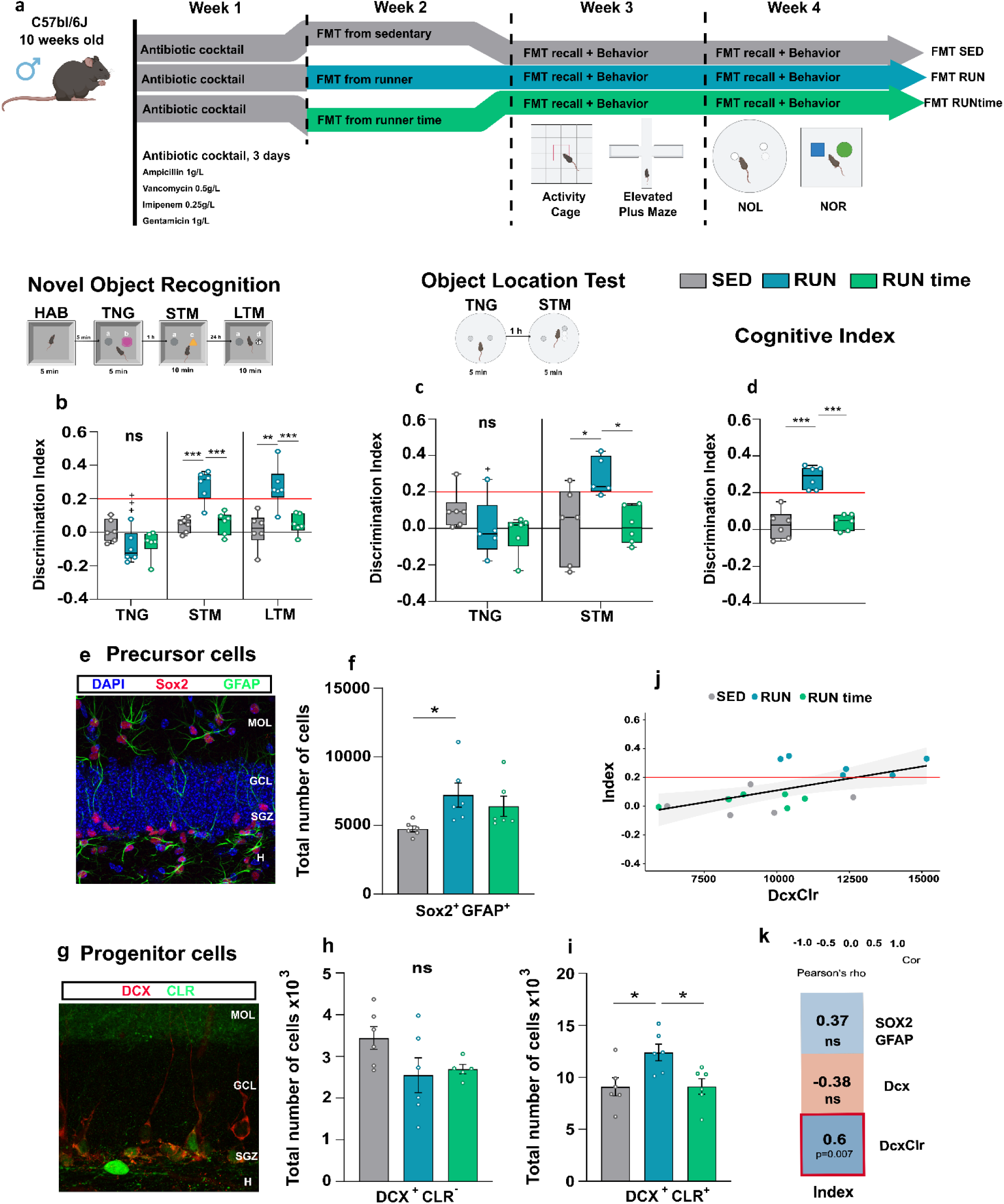
Fecal microbiota transplant from moderate runners induces cognitive improvement and promotes neurogenesis. **(a) Experimental design. (b)** Novel Object Recognition. Discrimination index in the three phases of the test. Linear model, F (8,45) = 14.1, p-value= 5.190e-10. One-way ANOVA is performed in each phase. (TNG: F (2.43), p-value=0.122; STM: F (21.8), p-value=3.65e-4, η2= 0.744; LTM: F (11.5), p-value=9.4e-4, η2= 0.605. **(c)** Novel Object Location. Discrimination index of the position of the columns in both phases of the test. Linear model, F (5,45) = 3.786, p-value= 9.59e-3. One-way ANOVA is performed in each phase. (TNG: F (1.75), p-value=0.21; STM: F (5.88), p-value=0.014, η2= 0.456). **(d)** Cognitive Index. One-way ANOVA, (F (31.852), p-value=3.99e-6, η2= 0.809). Intra-group comparisons, paired t-test or Wilcoxon test. **(e)** Representative image of double IF of SOX2 (red) and GFAP (green); DAPI staining is shown in blue. **(f)** Total number of SOX2+/GFAP+ cells in the SGZ, Kruskal-Wallis,(H _(_2,18) = 9.27, p-value = 0.009). **(g)** Representative image of double IF of DCX (red) and CLR (green). **(h)** Total number of DCX^+^/CLR^−^ cells in the GCL, one-way ANOVA, (F (2.133), p-valor=0.153). **(i)** Total number of DCX^+^/CLR^+^ cells in the GCL, one-way ANOVA, (F (5.366), p-valor=0.01, η2=0.433). **(j)** Correlation of the DCX^+^/CLR^+^ population with the CI (r = 0.6, p-value=0.007). **(k)** Correlations of NHA parameters with the CI. Red box represent significant correlation. Results represent mean ± SEM (n=6, SED group: n=6, RUN group; n=6, RUNtime group; n=6, RUNvel group). Between-group comparisons p-value *<0.05, **<0.01, ***<0.001. Ns, not significant. Intra-group comparisons p-value +<0.05, ++<0.01, +++<0.001.

No differences between groups were observed in motor activity parameters or anxiety-related behavior (not shown). The mice submitted to a FMT from RUN showed memory facilitation, both at the STM (F (21.8), p-value<0.0004, η2= 0.744) and LTM (LTM: F (11.5), p-value<0.001) in the NOR test, compared to animals receiving FMT from SED or RUNtime mice (Fig. 5b). We did not find differences in the exploration time at any of the test phases. Similarly, only the FMT RUN group had increased DI in the NOL test (STM: F (5.88), p-value=0.014, η2= 0.456; Fig. 5c) and in the CI (F (31.852), p-value<0.000004, η2= 0.809); Fig. 5d). Therefore, we have here demonstrated that the features of the microbiota of RUN mice confer cognitive facilitation to otherwise sedentary mice lacking that advantage.

Next, we examined the AHN variables of those mice submitted to FMT. We found increased SOX2^+^/GFAP^+^ (H (2,18) = 9.27, p-value<0.01; Fig. 5 e,f) and DCX^+^/CLR^+^ (F (5.366), p-value=0.01, η2=0.433; Fig. 5g,i) cells in recipients of FMT from RUN group compared to those receiving FMT from SED and/or RUN time donors, while no differences between groups were observed in DCX^+^/CLR^−^ cells (Fig. 5h). Again we revealed a significant positive correlation between the CI of the mice with their hippocampal DCX^+^/CLR^+^ cells (Fig. 5j,k). In summary, following the FMT experiments the proneurogenic effects of exercise were demonstrated in FMT recipients, replicating the results of their donor mice.

## Methods

### Animals

The animals used in the study were 82 male mice (Mus musculus L.) of the C57BL/6J strain (Envigo Laboratories), aged 11 weeks at the start of the experiments. They were all kept under hygienic conditions with a 12-hour light/dark cycle, stable temperature (20-22°C), and food and water *ad libitum* in accordance with the current European regulations (2010/63/EU).

All experiments were performed according to the European Community Guidelines (Directive 2010/63/EU) and Spanish Guidelines (Royal Decree 53/2013) and related rules, and they were first validated by the Committee of Ethics and Animal Experimentation of the Cajal Institute (20/05/2016), subsequently favorably evaluated by the CSIC Ethics Committee (Subcommittee of Ethics) of the Spanish National Research Council (07/27/2016) and authorized by the competent authority, the Animal Protection Area of the Department of Environment of the Community of Madrid (10/26/2016 and 06/19/2020).

### Study Design

All experiments involve a 32-days exercise protocol (6 weeks + 2 days) or home cage control. On the final day of the protocol, some behavioral test (Activity cage, EpM, NOL, or NOR) was conducted, and on day + 44, animals were sacrificed. All animals were randomly assigned to one of the following groups: SED: sedentary, RUN: moderate runner, RUNtime: intense time runners, RUNvel: intense velocity runners). All animals are 11 weeks old at the start of the experiment and are euthanized at 17 weeks of age.

### Treadmill running protocols

Every day of the treatment, the animals spent at least 30 minutes acclimating to the animal facility room where the treadmill was located before starting the exercise protocol.

On the first day, all groups were acclimated to the treadmill. Animals were allowed to explore the treadmill for 10 minutes without speed, and then the speed was gradually increased: after 2 minutes, it reached 300 cm/minute, after 5 minutes, 600 cm/minute, and finally, 10 minutes at the maximum speed of 1200 cm/minute. After those 10 minutes, the treadmill was gradually slowed down, maintaining a speed of 600 cm/minute for 30 seconds.

Starting from the second day, the SED group was not reintroduced to the treadmill. However, the sedentary group’s cage remained in the same animal facility room where the runners were performing their exercise task to ensure all groups were under the same conditions of odors, noises, and movements from one animal facility room to another. The exercise protocol followed by the RUN group for the remaining 6 weeks of the experiment was 1 minute at 600 cm/min, 2 minutes at 900 cm/min, and 37 minutes at 1200 cm/min. The RUNtime group followed the same gradual speed increase over the same time frame, and once reaching 1200 cm/min, they ran for 57 minutes. The RUNvel group ran for 40 minutes. During the first 3 minutes, the speed was gradually increased as in the other groups. However, in this protocol, the peculiarity was that the speed was continuously increased in short intervals until reaching 1800 cm/min, which was the maximum speed at which they ran for 3 minutes. Each group, after completing the prescribed time at maximum speed, the speed was reduced to 600 cm/minute, and after 30 seconds, the animals were returned to their cage without stopping the treadmill.

### Behavioral Assessment

All behavioral tests were performed during the light phase between 9 am and 3 pm (lights on from 7 to 7). All the tests were either recorded in video or analyzed automatically by the test device. When the test score was obtained by analyzing the videos, the experimenter was blind to the experimental groups.

#### Activity cage

This test was conducted using a Cibertec Actimeter XYZ-8 (MUX_XYZ16L-8 software) to assess locomotor activity. Animals underwent a two-day protocol, spending 5 minutes in the activity cage each day. On the first day, their spontaneous locomotor activity was measured for 5 minutes in a novel arena. The next day, the animals were placed in the same arena for another 5 minutes. The behavioral measures included total horizontal activity, total vertical activity, and total distance moved.

#### Elevated Plus Maze (EpM)

This test was conducted to assess whether any of the physical exercise protocols altered the anxiety state of the animals. The maze consists of two open arms and two closed arms, all elevated at a height of 0.5 meters above the floor. The task was performed in a single 5-minute session for each animal. Each animal was individually placed in the center of the maze, with its head facing towards the upper open arm (opposite to the experimenter). The animals were allowed to move freely around the maze for 5 minutes. The test was recorded on video for subsequent manual analysis by an expert experimenter. The following variables were taken into account: total time in each arm, total frequency in each arm, reaching.

#### Novel Object Recognition (NOR)

It was conducted using a challenging protocol within an open box measuring 42 x 32 x 31 cm. The challenging protocol of the test followed a procedure previously described in the work of McGreevy et al. (2019) and comprised four phases: habituation (HAB), training (TNG), short-term memory evaluation (STM), and long-term memory evaluation (LTM).

During the habituation phase, the animals were allowed to freely explore the empty box for 5 minutes. Immediately following, in the training phase, each animal was placed in the arena with two different objects in the center (objects A and B) and given 5 minutes to freely explore them. Subsequently, each animal was removed and returned to their home cage. One hour after training, animals were reintroduced to the NOR arena, which now contained a familiar object (object A) and a novel object (object C). They were allowed to explore these objects for 10 minutes. At the end of this phase, all animals were returned to their respective home cages in the animal room area. Twenty-four hours after the initial training, the animals underwent a 10-minute test in the arena, which still held the familiar object (A) and another new novel object (object D). The box was cleaned with a 0.03% acetic acid solution between trials.

The time spent exploring each object was manually recorded from the video footage. Discrimination indexes were calculated using the following formula for the training phase (TR), and in the short-term memory (STM) and long-term memory (LTM) phases, the time exploring object B was replaced by either object C or object D, respectively:

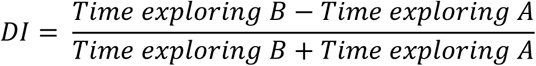

This formula quantifies the preference for exploring the novel object over the familiar object. A positive DI indicates a preference for the novel object, suggesting recognition or memory of the familiar object, while a negative DI suggests a preference for the familiar object. Between-group comparisons were conducted using a specific criterion for proper novel object discrimination, this criterion involves two key aspects:

1. Group Mean DI Scores: Each group must have a mean Discrimination Index (DI) score greater than 0.20. In other words, the groups should exhibit at least a preference for the novel object, with a bias of 60% exploration toward the novel object and 40% toward the known object.
2. Intragroup Difference: There must be a significant intragroup difference when comparing the DI scores for the short-term memory (STM) or long-term memory (LTM) phase to those of the training phase. This indicates that the preference for the novel object significantly increased in the memory phases compared to the training phase.

A criterion was set to establish a minimum exploration time, ensuring that animals that spent less than 1 second exploring each object were excluded from the analysis of the test results. In other words, animals that barely interacted with the objects were not considered in the analysis to maintain the reliability of the data.

#### Novel Object Location Test (NOL)

This protocol was conducted using a PVC circular arena measuring 35 cm in diameter and 20 cm in height, following the procedure described in (McGreevy et al., 2019). The test consisted of two phases: the training phase and the test phase. In the training phase, animals were allowed to explore the circular arena for 5 minutes. Two identical plastic columns were symmetrically placed along the diameter of the arena at the same distance from the walls. After this phase, the animals were returned to their home cages. One hour after the beginning of the training phase, the animals were reintroduced to the circular arena for 5 minutes (test phase). In this phase, one of the columns was displaced a short distance from its original position, creating a more challenging version of the test. The arena was cleaned with a 0.03% acetic acid solution between trials.

The time spent exploring each column was manually recorded from the video footage. Discrimination indexes (DI) were calculated using a specific formula:

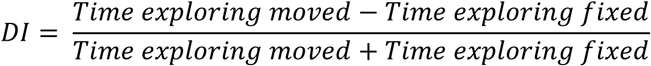

This formula quantifies the preference for exploring the displaced column relative to the non-displaced one and is used to assess spatial memory and object location recognition. In NOL, the column displacement distance determines the difficulty of the test. The shorter the displacement distance the more difficult the test. In this work we used only difficult versions of this test. Between-group comparisons were based on the same criteria mentioned above for NOR. A minimal exploration time criterion was used as explained above for NOR.

#### Cognitive Index (CI)

The Cognitive Index (CI) was calculated as a summary measure to evaluate cognitive ability and to compare the overall performance in various types of tests, including spatial and non-spatial tests, in both short-term and long-term trials conducted with different experimental groups. The CI was calculated as the mean (average) of the Discrimination Index (DI) scores from the three phases:

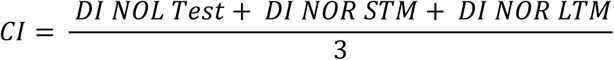

By calculating the mean of the DI scores from these three phases, the Cognitive Index provides a comprehensive overview of the cognitive performance of the experimental groups across different cognitive domains and timeframes. This single index allows for an easy and convenient comparison of overall cognitive abilities between groups.

### FMT protocol

#### Preparation of the Inoculum as FMT Treatment

Feces were collected from all experimental groups over three consecutive days. The animals were placed in a pre-sterilized compartment, and all the feces from each group were collected into a single 15 mL Falcon tube. The inoculum was prepared daily as the sample needed to be processed immediately. The entire process was carried out in a laminar flow hood. Pre-autoclaved PBS-Glycerol (1 mL of PBS per 100 mg of sample) was added to the collected feces. The mixture was then vigorously mixed to homogenize the sample. Subsequently, the solution was filtered using a 70 µm filter (ThermoFisher; REF: 15370801). Filtered solutions were frozen at −80°C for optimal preservation.

#### Procedure for FMT via Enema

Before proceeding with the FMT, the animals were treated with a mixture of antibiotics (Ampicillin 1g/L; Gentamicin 1g/L; Vancomycin 0.5g/L and Imipenem 0.25g/L) in their water for 4 full days, changing the water bottle each day and preparing a fresh antibiotic mix. On the fifth day, the antibiotic treatment was stopped, and the first FMT was administered. Animals were randomly divided into 3 groups, with each cage receiving FMT treatment from a different group: a) sedentary mice treated with FMT from sedentary mice (control); b) sedentary mice treated with FMT from the RUN group; c) sedentary mice treated with FMT from the RUNtime group. Before proceeding with the FMT, the animals were treated with a mixture of antibiotics (Ampicillin 1g/L; Gentamicin 1g/L; Vancomycin 0.5g/L and Imipenem 0.25g/L) in their water for 4 full days, changing the water bottle each day and preparing a fresh antibiotic mix. On the fifth day, the antibiotic treatment was stopped, and the first FMT was administered.

The FMT was performed via enemas under isoflurane anesthesia (IsoFlo, Zoetis Inc). Anesthesia was induced at 4% in a chamber, and once the mice were sedated, it was maintained at 1.5-2% during the procedure. Following the methodology of Zhou et al., (2019), once in the correct position (lateral decubitus), a plastic tube (2 mm inner diameter) connected to a 1 mL syringe was inserted into the colon through the anus of the mice to a depth of approximately 4 cm (tube length), 200 µL of the fecal bacteria suspension was drawn from the inoculum. After applying paraffin oil to the tube’s surface, the fecal bacteria suspension was slowly injected. The tube was then removed, and the anus was gently pressed with a cotton swab. The mice were kept in a prone position for 1 minute before being returned to their cages. Each animal received 5 consecutive FMTs, one per day, followed by 2 booster doses administered once a week. After the final booster dose, the mice were given a week of rest before performing cognitive tasks.

### Tissue collection

For the extraction of the mice’s brains, laboratory personnel anesthetized the animals with an intraperitoneal injection of sodium pentobarbital (Dolethal, Vetoquinol, France; 10 milligrams/kilogram), and limb reflexes were checked. A vertical incision was made in the abdomen to perform an intracardiac perfusion with 0.9% saline solution and extract the brain. Once this was done, the brain was divided down the midline into the left hemisphere, which was fixed by immersion in 4% paraformaldehyde (PFA) and the right hemisphere was dissected.

### Histology

Serial coronal brain sections with a thickness of 50 μm, including the hippocampal formation, were obtained from one hemisphere. This process was carried out using a Leica VT1000S vibratome. Each brain section was individually collected in a 96-multiwell plate filled with PB 0.1M. These plates were then stored at 4°C until they were ready for further analysis or examination.

#### Immunofluorescence

Each series of sections used for immunohistochemical analysis consisted in systematic sampling of the hippocampus in the rostro-caudal axis (8-9 hippocampal sections per animal, 50μm of thickness each section, 400μm apart from each other). For each immunofluorescence, the sections were kept floating with agitation in PBT-BSA blocking solution (PB with 1% Triton X-100 and 1% BSA) for five minutes. Primary antibodies (see table 1) were incubated in agitation with PBT-BSA for 1h at room temperature and 72h at 4°C. Secondary antibodies (see table 2) were incubated with PBT-BSA for 1h at room temperature and 24h at 4°C. Cell nuclei were counter-stained with 4’,6-diamino-2-phenylindole (DAPI, Sigma-Aldrich, 1:1000). For the DAPI staining, a random row was selected from the 96-well plate for each of the animals, and the sections were incubated for 12 minutes with DAPI at a 1:1000 dilution in 0.1M PB at room temperature with agitation. After 12 minutes, the sections underwent three washes with 0.1M PB and were mounted on slides, arranged from rostral to caudal, and secured with gelvatol beneath a coverslip.

**Table 1.** List of primary antibodies for IF.

| Antibody | Manufacturer | Cat number | Dilution |
| --- | --- | --- | --- |
| **CLR** | Santa Cruz | sc-8066 | 1:500 |
| **CLR** | Synaptic Systems | 214 104 | 1:500 |
| **DCX** | Swant | 7699/4 | 1:3000 |
| **DCX** | Abcam | ab18723 | 1:1000 |
| **GFAP** | Abcam | Ab7260 | 1:1000 |
| **GFAP** | Sigma Aldrich | G3893 | 1:1000 |
| **SOX2** | R&D | af2018 | 1:200 |
| **pH3** | Millipore | 06-570 | 1:500 |
| **Lectin** | Sigma Aldrich | L0401 | 1:100 |
| **PDCX** | R&D Systems (Biotechne) | AF1556 | 1:1000 |
| **ZO1** | Thermo Fisher Scientific | 61-7300 | 1:500 |
| **Iba1** | Wako - Fujifilm | 019-19741 | 1:1000 |

**Table 2.** List of secondary antibodies for IF.

| Antibody | Manufacturer | Cat number | Dilution |
| --- | --- | --- | --- |
| **Alexa anti-Rabbit 488 nm** | Santa Cruz | sc-8066 | 1:1000 |
| **Alexa anti-Rabbit 594 nm** | Synaptic Systems | 214 104 | 1:1000 |
| **Alexa anti-Rabbit 680 nm** | Swant | 7699/4 | 1:1000 |
| **Alexa anti-Mouse 594 nm** | Abcam | ab18723 | 1:1000 |
| **Alexa anti-Rat 488 nm** | Abcam | Ab7260 | 1:1000 |
| **Alexa anti-Goat 594 nm** | Sigma Aldrich | G3893 | 1:1000 |
| **Alexa anti-Goat 488 nm** | R&D | af2018 | 1:1000 |

First, IF of adult hippocampal neurogenesis (AHN) markers was performed, as the positive effects of exercise on learning and memory are mediated, among other factors, by AHN. For this, a double staining of SOX2/GFAP was conducted to identify neural precursors. Additionally, histone 3 phosphorylated (pH3), an intrinsic proliferation marker detected during the mitosis phase was performed to analyze the rate of proliferative cells in the DG of the hippocampus. Double staining of doublecortin (DCX) and calretinin (CLR) was also carried out. This double labeling allowed the study of different populations: DCX^+^/CLR^−^ cells identify type 2b neural progenitors (highly proliferative cells) and type 3 cells (cells in transition from a potentially proliferative state to an immature post-mitotic neuron); DCX^+^ and CLR^+^ cells correspond to immature neurons in differentiation, i.e., post-mitotic cells; finally, DCX^−^/CLR^+^ cells correspond to immature neurons in later stages of differentiation, a short-lived phase that gives rise to mature granular neurons.

For the study of barriers, we used different markers; for blood vessels, we used podocalyxin (PDCX) and tomato lectin. Podocalyxin is a protein found in endothelial cells of blood vessels, while lectin is a substance that specifically binds to carbohydrates present on the surface of endothelial cells, allowing specific visualization of blood vessels. To mark tight junctions, we used zonula occludens-1 (ZO1) protein (a component of the tight junctions between endothelial cells). For microglia, we used ionized calcium-binding adapter molecule 1 (Iba1) and glial fibrillary acidic protein (GFAP) for astrocytes.

### Image Analysis of Fluorescence Microscopy

#### Cavalieri method

The volumetric analysis of hippocampal structures, specifically the granular cell layer (GCL) and the length of the subgranular zone (SGZ) in the dentate gyrus (DG) of the hippocampus, was conducted using the Cavalieri method. This stereological method estimates the volume of a reference space by summing the areas of a fraction of its sections and multiplying by the constant distance between them.

For this analysis, sections stained with Nissl or DAPI were used. The THUNDER microscope (Leica Microsystems) was employed to obtain between 6 and 7 images per animal of the hippocampal dentate gyrus. Subsequently, ImageJ software was used to quantify the surface area of the SGZ and the area of the GCL in each image. To measure the GCL area, the “polygon tool” in ImageJ was used, while the “segmented line tool” was used to measure the SGZ length. Before taking the measurements, the scale of each image was set. This process enabled precise measurements of the volume of the hippocampal structures of interest in each analyzed animal.

To calculate the volume of the GCL in the dentate gyrus of the hippocampus using the Cavalieri method, the sum of the GCL area (A) of each image for each animal was multiplied by the known distance (d) between the images. Given that the coronal sections were 50μm thick and a row of sections from a 96-well plate was used, the distance between sections is 400μm (considering there is one section every 8 wells in a 96-well plate). Thus, the volume of the GCL is estimated using the formula:

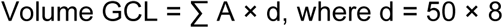

By multiplying the area of each image by the distance between sections and summing the results for all images, an estimate of the total volume of the GCL in each animal is obtained. This method provides a precise volumetric measure of the structure of interest in the hippocampus. To calculate the area of the SGZ in the dentate gyrus of the hippocampus, a modification of the Cavalieri Principle was used. Instead of multiplying the area by the distance to find the volume, since the aim is to find the area, the length (L) of the SGZ in each image was measured and multiplied by the distance.

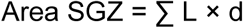

### Stereology

#### Fractionator Method

This method was used to count PH3^+^ in the SGZ of the dentate gyrus. It involves manually counting the labeled or positive cells using a fluorescence optical microscope (Leica DMI6000B) with a 40x objective. First, the state of the hippocampus in all sections of the animal is checked at 10x magnification to ensure the dentate gyrus is intact and no section is missing from the series; if any damage or missing section is found, the animal is immediately discarded from the analysis. Next, the objective is switched to 40x to examine the entire subgranular zone of the dentate gyrus, counting only the positive cells present in that area. This count is conducted in one of the 8 series obtained by cutting with the vibratome, so the total number of cells counted in each animal corresponds to the number of proliferative cells in a fraction of the hippocampus, hence the name, fractionator. Specifically, staining and counting were performed in 1/8 of the hippocampus of each animal. Therefore, to calculate the total number of positive cells in the hippocampus, this fraction is multiplied by the number of series, 8 series.

#### Disector Method

In the case of markers like DCX/CLR and SOX2/GFAP, the fractionator method cannot be applied due to the large number of immature cells present in the dentate gyrus. Instead, we applied a physical dissector in U, following the method previously described in the laboratory (Llorens-Martín et al., 2006).

For <u>DCX/CLR</u>, approximately 6 blocks of 11 images each were taken using a direct confocal microscope (Leica TCS SP5) with a 63x immersion objective, 2.5x zoom, 1.72 µm step size, and 1024 x 1024 pixels. Images were captured from different sections of the GCL and SGZ, randomly selecting 1 block of images from the rostral zone, 2 from the medial zone, and 3 from the caudal zone. Counting was performed using a simple blind count with ImageJ. All images in each block were analyzed, moving along the z-axis, evaluating the 594 nm channel (red, marking DCX), the 488 nm channel (green, marking CLR), and finally both channels together. This allowed us to distinguish between different cell types: DCX+/CLR- and DCX+/CLR+. The criteria applied to consider a cell positive were as follows: None of the analyzed cell types should touch the exclusion borders, which are the left and lower edges of the photograph, in addition to the last photo of each block; The marker had to be present in at least 3 of the images in the block. This means that if a cell had a marker in only 2 photos of the set or block but not in the third, it was not considered positive for that marker

In the case of <u>SOX2/GFAP</u>, 7 blocks were used, each consisting of approximately 15 images. A direct confocal microscope (Leica TCS SP5) was used again, with a 40x objective, 2.4x zoom, 2.01 µm step size, and 1024 x 1024 pixels. Images were captured from different sections of the GCL and SGZ of the dentate gyrus in the hippocampus, randomly selecting 1 block of images from the rostral zone, 2 from the medial zone, and 4 from the caudal zone. We conducted the counting using a simple blind count with ImageJ from Fiji. For this, a digital grid was placed over the image, and each block was analyzed by moving along the z-axis. The criteria used to consider a cell positive as follows: The soma of the cell is located in the subgranular layer of the dentate gyrus. The SGZ is defined as a distance of about 3 somas from the inner boundary of the granular layer; the cell has a single apical dendrite that may branch distally from the soma, but never at the base; the soma of the cell does not touch any exclusion borders (left edge of the photograph and last photo of the block). Once the total number of cells of each population was obtained, we estimated the cell density for each block of images. After calculating the cell density for each block, this allowed us to extrapolate the total number of cells for each individual with the area of SGZ and the volume of GCL.

#### Analysis of Vascularization by Immunofluorescence

For vascularization analysis, we performed Lectin immunofluorescence and estimated the area occupied by fluorescence for each image. Additionally, we used the AngioTool software, following the methodology of (Zudaire et al., 2011). AngioTool is designed to automatically analyze vessel images, allowing for the quantification of various parameters such as vascular density, vessel length and diameter, vascular branching, and more, providing a broader view of the structural complexity of the vascularity in our study area.

To perform this analysis, we captured images using a STELLARIS8 confocal microscope, to create mosaics of the entire hippocampal formation, taking 3 blocks from each animal. These blocks were randomly selected: 1 rostral block, 1 medial block, and 1 caudal block. We used a 20x objective, with a resolution of 2048 x 2048 pixels, a step size of 1.01 µm, and a zoom of 0.75. Additionally, we captured images of the entorhinal cortex, which we used as a reference to determine whether our exercise protocols specifically affected the hippocampus or also other regions. For quantification, we decided to generate 3 crops in each image within the hippocampal formation. We selected one crop in the hilus, another in the molecular layer, and another near the CA1 area.

#### Quantification of tight junctions

For ZO1, we selected 3 hippocampal sections: one rostral, one medial, and one caudal. Within each section, 3 photographs were taken in 3 different zones, where a main vessel was identified. The photographs were captured under the following conditions: using a 63x objective, with a resolution of 1024 x 1024 pixels, and a step size of 1.01 µm. For quantification, we used the projection of the images from each block. We separated the channels to work with the vessels (594nm channel, podocalyxin) on one side and the tight junctions on the other (488nm channel, Zo1). After separating the channels, we applied the automatic thresholding method OTSU for both vessels and tight junctions, and measured the area occupied by each channel. Additionally, we selected a vessel of interest using the magic wand tool and added it to the ROI manager, which we then used to implement this ROI in the tight junction channel to measure the percentage of tight junction area within a vessel.

#### Analysis of Glial Cells (Astrocytes and Microglia)

For the quantification of both astrocytes and microglia, we used a double staining of GFAP and Iba1 along with Dapi for the localization of the Granule Cell Layer (GCL) in the Dentate Gyrus. We captured a block of approximately 5 images in each hippocampal section. In each photo, we took a cell as a reference and captured the entire thickness in which that cell was located under the following conditions: using a 40x objective, with a resolution of 1024 x 1024 pixels, and a step size of 2.01 µm.

Once the photos were taken, we quantified the occupied area and the fractal dimension of each zone of the hippocampal formation, Hilus (H), Granule Cell Layer (GCL), and Molecular Layer (ML). With the DAPI staining, we assisted in defining the GCL, to be able to count within this layer or in the rest of the layers. After selecting the regions, we worked on the image projection, we applied an automatic threshold to generate a binary image. After testing all the methods offered by ImageJ, we settled on “triangle”, which best fit our signal in both GFAP and Iba1. On this image (projection + “triangle” threshold), we examined two things: structural complexity and signal occupied, also known in ImageJ as “Area” and “Area fraction”.

Once we obtained the measurement of the signal-occupied area, we proceeded to measure the structural complexity. For this, we used a mathematical concept called fractals; in summary, the higher the fractal dimension, the greater the structural complexity. We performed this measurement using a “fractal box count” tool, which provided us with the fractal dimension of our image. We analyzed these two variables in both GFAP and Iba1, but in the case of Iba1, we also counted the cells. For this, we used “analyze particles”, which removed all individual signals that did not meet certain criteria. In our case, the conditions set were to remove anything smaller than 70 pixels, to leave the soma intact and facilitate the counting of microglial cells. We counted these cells as cell profiles, as we did not reference them to a total area or volume since it was the same in all animals.

### 16S Amplicon Microbiota composition

#### DNA Extraction and 16S Library from the Cecum

DNA extraction was performed using the QiaAMP Power Fecal Pro kit (QIAGEN) following the manufacturer’s instructions. DNA concentration was normalized, and 16S rRNA amplicon libraries were prepared using primers to amplify the V3-V4 region of the bacterial 16S rRNA gene, with Illumina adapters incorporated as described in the Illumina 16S rRNA metagenomic library preparation guide, except using 30 amplification cycles. After index PCR and purification, the products were quantified using the Qubit High Sensitivity DNA kit (Life Technologies) and pooled equimolarly.

Pooled libraries were evaluated using an Agilent High Sensitivity DNA kit and examined by quantitative PCR (qPCR) using the Kapa quantification kit for Illumina (Kapa Biosystems) following the manufacturer’s guidelines. Subsequently, libraries were diluted, denatured following Illumina guidelines, and sequenced (2×300 bp) on the Illumina MiSeq platform.

#### Bioinformatic Analysis of Cecum Microbiota in Mice

All bioinformatic analysis was conducted using the R programming language (version 4.3.3) through the RStudio interface (version 2023.12.1+402 “Ocean Storm”). Raw sequencing files (forward and reverse) and a metadata file containing information about which animal each sample belongs to and behavioral data of each animal were loaded. The number of reads in the raw files ranged from 34,002 to 83,327. After loading the data, a quality plot of the reads was generated to establish cutoff points for each sequence. Using the R DADA2 library, with the trim and filter function, 37 bp were trimmed from the left and 50 bp from the right (cutoff point at base pair 250). Error learning was computed from our data to perform denoising, where sequences classified as noise were removed, and finally, forward and reverse reads were merged, and chimeras were removed. The total number of reads retained for analysis after filtering ranged from 20,310 to 47,430. In terms of the percentage of unfiltered reads, they ranged from 56.3% to 59.9%. No samples were eliminated due to a low number of reads, either initially or after filtering.

Unfiltered reads were taxonomically assigned using the SILVA v138.1 database. Species were also assigned using the SILVA species database, v138.1. Sequences were aligned using the ClustalW algorithm through the R package “msa” to compute the tree and phylogenetic distances between sequences (optimized using the “GTR” model). Both the data from the reads (amplicon sequence variants or ASVs), taxonomy, and the phylogenetic tree were combined into a phyloseq file using the package of the same name to work on it. First, taxonomic filtering was performed at the genus level. All sequences with unknown genera were removed, leaving 96 genera. Next, the percentage composition per sample was graphed (using the ggplot2 library), grouping all genera that accounted for <1% of the total of each sample under the label “Rare Taxa”.

Alpha diversity was calculated using the “Tjazi” package, specifically the Chao1, Simpson, and Shannon indexes ^48^. To check for differences between groups, a one-way ANOVA was computed for each index and subsequent pairwise comparisons (with p corrected by Tukey’s HSD) when significant. Graphs were also made using ggplot2. For the calculation of beta diversity, a new filtering was performed, removing any sequence that was not present in at least 10% of the samples (for our n=19, each sequence had to appear in at least 2 samples, rounding up). Eight genera were removed, leaving 88. Additionally, a CLR transformation (Centered Log-Ratio) was performed. Group differences were calculated using the “adonis2” library, using a PERMANOVA, with Euclidean distance and 1000 permutations. Pairwise comparisons were conducted using the “ecole” library with the “permanova_pairwise()” function. Graphically, it was represented using a PCA, also using ggplot2.

For differential abundance, the “fw_glm()” function of the “Tjazi” library was used, using the parameter “order = “atc””. Thus, we have the results of the model and all pairwise comparisons. With a cutoff of 0.1 in the FDR-adjusted p values of the models, 8 significantly different genera were found. When calculating the correlation of these 8 genera with the mean score obtained in the behavior tests, 4 significant correlations were found, all with ∣ρ∣ > 0.51.

### Statistical analysis

For comparisons between two independent groups, the T-test was applied when the dependent variable was normally distributed. The Mann-Whitney U Test was used in the case of non-normal distribution. For comparisons between two dependent groups, paired sample t-test was applied if the dependent variable was normally distributed. When not, Wilcoxon Signed Ranked Test was used. For behavioural tests that contained between-subject and within-subject measures with three levels, Mixed ANOVA was applied when met the normality and homogeneity assumptions. If the normality and/or the homogeneity of variances assumptions were violated, a Friedman test was used followed by a post hoc Wilcoxon Signed-Ranked Test. To study correlations, Pearson correlation was calculated when data followed normal distribution. Spearman correlation was used with non-normal distributions. All data were analyzed using SPSS Statistics (IBM, v.26.0.0). Data are shown as mean ± SEM. To test normality, the Shapiro-Wilk Test was applied. Extreme values identified by the software were eliminated from the analysis.

Inter-group differences: *p<0.05, **p<0.01, ***p<0.001. Trends in inter-group differences 0.05 ≥ *’p < 0.099. Within-group differences: +p<0.05, ++p<0.01, +++p<0.001. Trends in intra-group differences 0.05 ≥ +’p < 0.099. All graphs were created in GraphPad Prism 8.

## Supplementary Figures

**Sup.Fig. 1.**
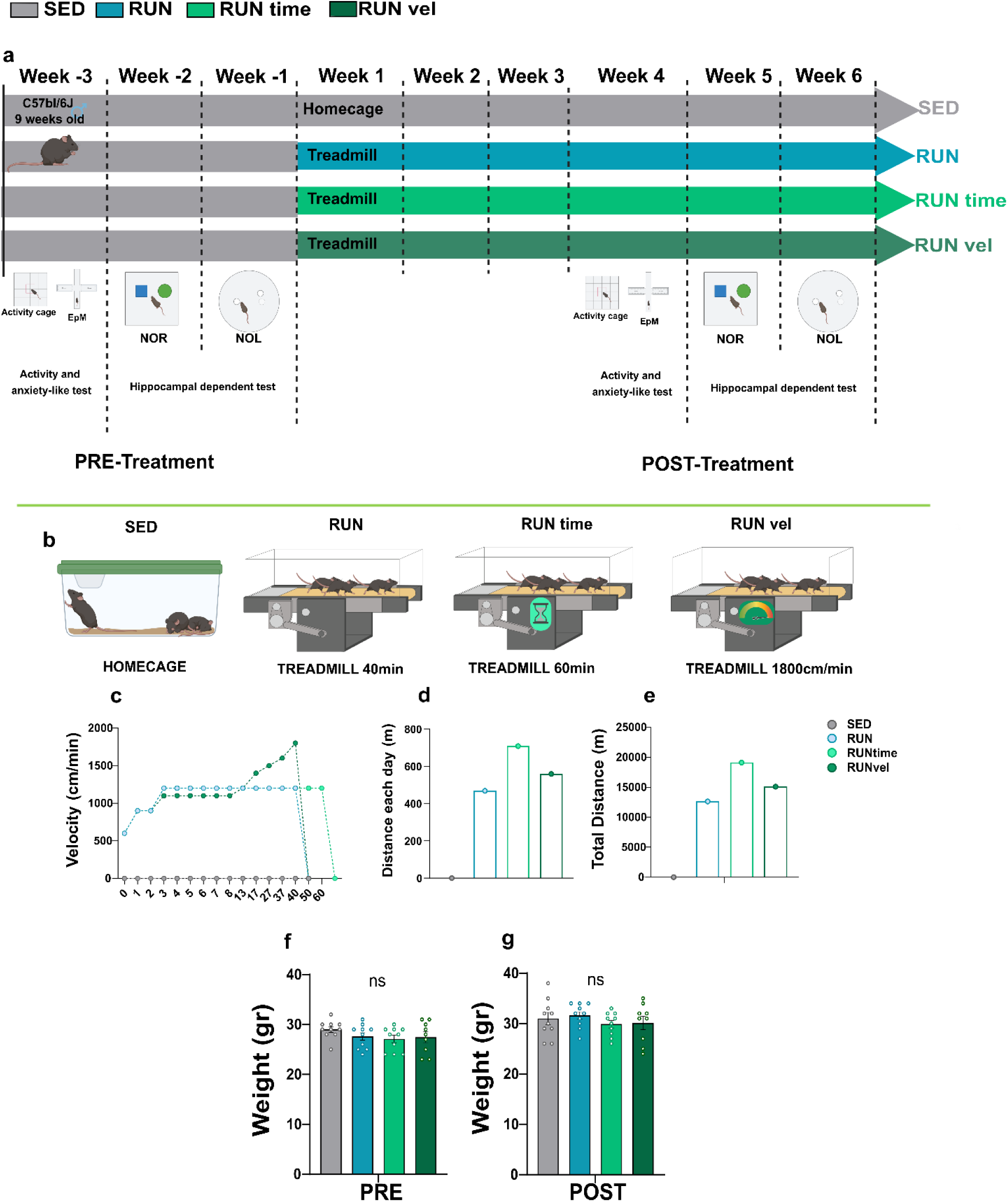
Experimental Design. (a) Scheme of the longitudinal and between-group design we followed. We conducted a series of behavioral tests both before treatment (pre-treatment) and after 6 weeks of physical exercise (post-treatment), using approximately 3 weeks to perform all behavioral tests. (b) Representation of the 4 experimental groups and the protocol for each group. (c-e) Representation of the characteristics of the physical exercise protocols. (c) Graph depicting the speed at which each group runs at each minute of the daily protocol. (d) Distance in meters covered by each group daily. (e) Distance in meters covered by each group in total after the completion of treatment (5 days per week, 6 weeks in total). (f-g) Weights in grams (f) before physical exercise treatment and (g) after physical exercise treatment.

**Sup.Fig. 2.**
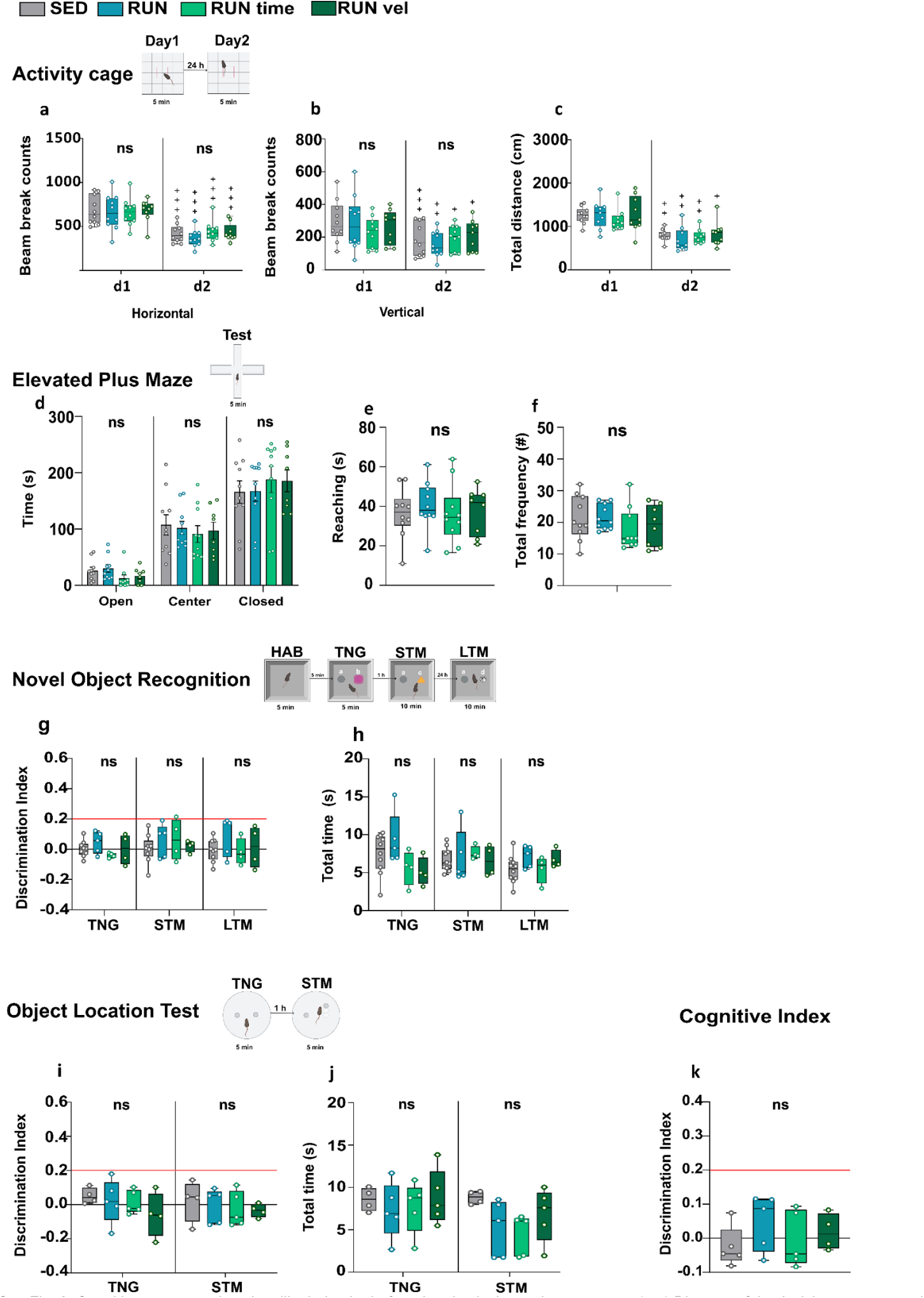
Cognitive, motor, and anxiety-like behavior before the physical exercise treatment. (a-c) Diagram of the Activity cage protocol and variables. (a) Total horizontal activity, one-way ANOVA for each day (Day 1, F (0.045), p-value = 0.987); (Day 2, F (1.03), p-value = 0.391). (b) Total vertical activity, one-way ANOVA for each day (Day 1, F (0.474), p-value = 0.702); (Day 2, F (0.64), p-value = 0.595). (c) Total distance traveled, Kruskall-wallis for each day (Day 1, H (3, n=39) = 2.37, p-value = 0.499); (Day 2, H (3, n=39) = 1.38, p-value = 0.711). (d-f) Diagram of the EPM protocol and variables. (d) Time in arms, Kruskall-wallis for the 3 phases. (Open; (H (3, n=38) = 7.87, p-value = 0.049), Center; (H (3, n=38) = 1.15, p-value = 0.765), Closed; (H (3, n=38) = 1.54, p-value = 0.674). (e) Reaching, one-way ANOVA (F (0.246), p-value = 0.863). (f) Total frequency, Kruskall-wallis (H (3, n=38) = 3.65, p-value = 0.301). (g-h) Diagram of the NOR protocol and variables. (g) Discrimination Index in the three phases of the test. Mixed ANOVA. Group effect (F (3,18) = 0,410, p-value = 0,748), phase effect (F (2,36) = 2.495, p-value = 0,097). One-way ANOVA in each phase. (TNG: F (1.32), p-value = 0.9; STM: F (0.512), p-value = 0.9; LTM: F (1.32), p-value = 0.9). (h) Total time exploration of both objects in the three phases of the test. Group differences in each phase, one-way ANOVA. (TNG: F (2.40), p-value = 0.3; STM: F (0.155), p-value = 1; LTM: F (0.291), p-vaue = 1). (i-j) Diagram of the NOL protocol and variables. (i) Discrimination Index of the column position in both phases of the test. Mixed ANOVA. Group effect (F (3,14) = 1.029, p-value = 0.410), phase effect (F (1,14) = 0.259, p-value 0.618). One-way ANOVA in each phase. (TNG: F (0.871), p-value = 0.958; STM: F (0.825), p-value = 0.958). (j) Total time exploration of both column objects in each phase of the test. Group differences in each phase, one-way ANOVA. (TNG: F (0.278), p-value = 0.84; STM: H (3, n=20) =6.57, p-value = 0.087). (k) Cognitive Index. One-way ANOVA, (F (0.942), p-value = 0.445). Results represent the mean ± SEM (n=10, SED group: n=10, RUN group; n=10, RUNtime group; n=9, RUNvel group). Group comparisons p-value *<0.05, **<0.01, ***<0.001. Ns, not significant. Intra-group comparisons p-value +<0.05, ++<0.01, +++<0.001.

## Discussion

In the present work, we investigate the impact of different exercise intensities and durations on brain function and cognition, highlighting the potential mediating role of the gut microbiota. Our findings support the hormetic effects of exercise, demonstrating that moderate exercise (40 minutes daily at 1200 cm/min) enhances cognitive function and adult neurogenesis compared to sedentary controls. However, these benefits are lost with longer or more intense exercise, illustrating the hormetic profile of exercise effects on the brain. Additionally, we observed variations in both alpha and beta diversity in the gut microbiota, and the microbiota composition profiles closely related to the volume of physical activity performed, ranging from sedentary to high-intensity or long-duration exercise.

Notably, the relative abundance of certain bacteria correlated with the cognitive performance of the different groups. Through fecal microbiota transplant (FMT) experiments, we further established a possible causal relationship, showing that the cognitive and neurogenesis benefits observed in runner donors could be transferred to sedentary animals. These findings highlight the role of the gut microbiota in the cognitive effects of moderate exercise, suggesting it as a key mediator in the hormetic response.

Regarding the mechanisms that mediate the effects of physical exercise on the brain, a large amount of evidence has accumulated in the last two decades. These studies have focused on mediating factors both outside and inside the brain ^2,49–53^, including increased blood circulation at the vascular niche ^54–56^ and AHN ^57,58^, as well as recent genetic analyses on exercise’s influence on brain function and memory ^59^. Although some of these mediating factors of the effect of exercise have been studied in the context of their hormetic profile ^6,8^, there is still very little information about the mechanisms supporting it.

A major challenge is the variability in defining “moderate” versus “strenuous” exercise across studies, and inconsistencies in the analyzed variables, contributing to debates on exercise’s effects on cognition in humans ^60–63^.Despite this, there is consensus that mechanisms underlying the hormetic response in mice should reflect the observed physiological outcomes, such as cognitive performance.

In the present work we focused on AHN and gut microbiota as potential mediators of exercise effects. We specifically tested whether commonly used exercise protocols (1 hour/day, 5-7 days a week, 1700-2200 cm/min, near or above the supra-lactate threshold; ^8^ would have a positive effect on memory, comparable to our moderate protocol (40 minutes/day, 5 days a week, 1200 cm/min). We have found that only the moderate exercise protocol improved recognition memory, consistent with previous studies by our group ^13,47^ and other laboratories ^12,64–66^, for review, see for example ^67,68^. To maximize the possible differences between distinct exercise protocols among them and with a sedentary lifestyle, we designed behavioral tests with difficulty thresholds that sedentary animals could not adequately reach. This approach demonstrated that only moderate exercise enhances cognitive performance, a novel finding in adult laboratory mice.

Although our experimental design did not include an exhaustive dose-response curve, our results allow us to describe three exercise protocols at different points of the hormetic curve: moderate exercise near the maximum stimulatory response, longer duration and higher speed protocols near the non-observable effect level (NOEL), and the sedentary group represents the zero end-point (ZEP) level ^69–71^.These points were also evident when analyzing potential mediating mechanisms, both within the brain (AHN) and outside (microbiota).

Our analysis shows that the moderate exercise significantly increased both the total number of neural precursors and immature neurons in the hippocampal granule cell layer, an effect not observed with more intense or prolonged protocols. These findings align with well-established role of AHN in cognition, learning, and memory ^72–74^ and support previous evidence that cognitive benefits of exercise are linked to increased AHN, particularly the number of immature neurons ^75–77^.

Recent studies have also highlighted the influence of the microbiome on AHN regulation in rodents ^29,78–83^. To explore whether the microbiota mediates exercise-induced changes in AHN and cognition, we analyzed the neurogenesis rate and performance in hippocampal-dependent tasks in animals receiving fecal microbiota transplants (FMT) from exercised donors. Our results demonstrated that cognitive improvements and increased AHN could be transferred to sedentary recipients via FMT, suggesting a microbiota-AHN pathway in exercise-induced cognitive effects.

Altogether, our present results suggest that one of the exercise-induced actions on cognition is through a microbiota-AHN pathway. Besides, our findings indicate that while forced intense exercise does not promote AHN nor cognitive enhancement, moderate exercise effectively does, consistent with prior research linking this outcome to increased lactate and corticosterone levels ^8,15^. Interestingly, studies comparing aerobic and resistance exercise protocols found that intense aerobic exercise enhanced neurogenesis without corresponding cognitive benefits ^84^. Additionally, microbiota changes induced by moderate exercise could potentially act through the vagus nerve ^29,85^, or through metabolites ^26,86^, both mechanisms warranting further investigation.

Regarding microbiota composition, our analysis aligns with previous research on aerobic exercise, particularly when comparing sedentary subjects. The literature on microbiota and exercise is often confounded by differences in study design, such as variations in mouse models, diet, and other lifestyle conditions. Despite these challenges, our findings are consistent with previous studies that suggest moderate exercise promotes greater microbial diversity. Several studies have demonstrated that differences in microbiota composition were noted based on the type of exercise, such as voluntary versus forced exercise ^87^ or strength versus endurance training ^88^. Notably, our study is the first to distinguish these differences using different intensities of the same type of exercise protocol, specifically distinguishing exercises based on their cognitive effects.

We found that only the moderate exercise protocol led to cognitive improvement, suggesting that this effect is specific to moderate exercise. Our data indicate that exercise modulates a significant number of bacterial genera (15 in total). We highlight four genera: *Angelakisella*, *Erysipelatoclostridium*, *Coriobacteriaceae UCG*-*002*, and *Acetatifactor* for their strong correlation with cognitive performance. Changes in *Lachnospiraceae* and *Oscillospiraceae* families, linked to cognition in previous studies, were observed exclusively in the moderate exercise group ^89,90^ and reductions in genera like *Prevotellaceae* UCG-001 and *Lachnospiraceae* UCG-066 were associated with poorer cognitive performance <u>(D’Amato et al., 2020</u>). Our findings are consistent with previous research, such as the increase in *Eubacterium ruminantium* observed across different types of exercise ^91^ and the reduction in *Alloprevotella*, similar to reductions reported for the entire *Prevotella* taxon ^92,93^.

We also found an increase in *Clostridium sensu stricto* (strain 1) only under exercise conditions of one hour, which aligns with several findings, such as chronic fatigue syndrome ^40^ and healthy individuals, both murine models ^87^and humans ^94^. In our data, Streptococcus appeared exclusively in the moderate exercise group, with several studies associating this genus with beneficial effects ^95^. We also observed an increase in several *Lachnospiraceae* species unique to the moderate exercise group, consistent with other findings linking this increase to improved memory^96^. Interestingly, in older mice, where cognition is compromised, voluntary exercise increased Alloprevotella while reducing Lachnospiraceae and Acetatifactor ^97^, suggesting that moderate exercise may offer specific cognitive benefits compared to sedentary or intense exercise.

Further studies also agree with our findings regarding exercise-induced microbiota changes, with specific variations depending on the taxon. For instance, we found a decrease in *Rikenellaceae* RC9 in high-intensity or long-duration exercise groups, with no changes between moderate and sedentary exercise, whereas ^98^ reported an increase in humans. Differences in study design, such as dietary supplementation (e.g., with *Streptococcus thermophilus*), may account for these discrepancies. However, this study agrees on an exercise-induced increase in *Oscillospiraceae*, which appeared exclusively in our moderate exercise group. ^99^ reported an increase in *Prevotella*, contrary to our data and others (see references above). This discrepancy may be due to differences in training duration, as longer training times have been associated with higher *Prevotella* levels ^100^. Variations in exercise protocols, data analysis methods, and mouse strains further complicate direct comparisons between studies.

In summary, alterations in microbiota composition appear to contribute to the cognitive benefits associated with moderate exercise, particularly through enhanced adult hippocampal neurogenesis (AHN).

Barriers are an indicator of the state of both the brain and the gut, as they are affected by variables such as oxidative stress, cytokines, growth factors, etc. ^101,102^. Although it is quite common in the literature to associate the decrease of some tight junction proteins with a state of barrier permeability, this assertion is delicate because is usually consider a more complex process ^103^. We found it interesting to observe what happens to certain tight junctions in both barriers (blood-brain and intestinal epithelium) with our exercise protocols, as it is common to relate physical exercise with alterations in barriers ^44,104,105^. We observed that tight junctions in the gut do not seem to fluctuate much, at least at the messenger level, while in the hippocampus, changes are dependent on the intensity of the exercise. We observed changes in Claudin 5 levels in the hippocampus, fluctuating in relation to exercise intensity. Additionally, ZO-1 levels seem to be elevated only in the moderate group. This increase could be an adaptive change, as ZO-1 is an essential protein necessary for tight junctions. It seems as if the moderate group is more prepared in case more junctions need to be formed. Regarding barriers, there are few studies showing fluctuations in these proteins merely due to physical exercise, and even fewer with protocols similar to ours. Regarding the blood-brain barrier, a recovery in ZO-1, occludin, and claudin 5 levels has also been observed thanks to exercise, both in APP/PS1 animals ^106^ and animals on a high-fat diet ^102^. An increase in BBB permeability may consequently allow serum glucocorticoids (or another factors) to enter the brain, potentially impacting AHN and impairing the production of brain-derived nerve growth factor (BDNF).

It is important to note that the present results cannot be explained as a consequence of high levels of exercise inducing no actions on the body in general and the brain in particular. In fact, the most intense exercise protocols do manifest specific effects at various levels, both outside, such as microbiota composition, already discussed, and within the brain, where we found microglial activation. Indeed, microglial area and complexity were increased in the mice exercised with the more intense protocols, but absent in the moderate group. Microglia are resident brain macrophages that become activated and proliferate following brain damage or stimulation by immune mediators. On the other hand, quiescent microglial cells sustain neuronal health by secreting different trophic factors. It is well known that voluntary exercise decreases microglial activation under neuroinflammatory conditions occurring in models of aging or neurodegenerative diseases, brain injury such ischemia or after endotoxin administration ^107^. Moreover, inflammatory blockade restores adult hippocampal neurogenesis ^108^. Conflicting results have been reported in regard microglia following physical exercise. Some authors have described increased proliferation of microglia in cortical regions ^109^, while others found no changes after a moderate protocol ^110,111^, and ^112^ reported that AHN was inversely correlated with microglial density. In summary, absence of effect of intense exercise on AHN might rely on microglial activation as well. On the other hand, microbiota impacts on microglia. Indeed, it is known that microbiota shapes microglial maturation and function ^113^. In addition, germ-free mice show microglial alterations compared with animals bearing normal gut microbiota ^114^. Therefore, we suggest that hippocampal microglial activation in intense runners in the present work involve exercise-induced microbiota alterations.

In conclusion, our results demonstrate that the beneficial effects of moderate exercise on specific learning and memory tasks in mice vanish with small increases in daily training time or increases in running intensity, resembling a hormetic curve profile for these cognitive effects of exercise. This profile is paralleled by the AHN and microbiota composition changes. Moreover, an exercise intensity-specific microbiota signature can be outlined, suggesting a potential mediator of the hormetic effects of physical exercise on brain and cognition. Furthermore, the cognitive benefits induced by fecal microbiota transplant from moderate exercised to sedentary animals are not observed when the transplants come from animals running for longer times, which is consistent with the same profile seen in AHN. Altogether, our results indicate that a microbiota-AHN pathway mediates the hormetic effects of physical exercise on the brain and cognition.

## ACKNOLEDGEMENTS

We are grateful to Laude Garmendia from the Animal House, at the Cajal Institute for her unpayable help and advice, to the Image Analysis Unit of the Cajal Institute, and to the Laboratory of Omic Technologies and Bioinformatics of the Cajal Institute. E.C. and P.M. were funded by predoctoral fellowship (FPI) grants from the Spanish Ministry of Economy and Competitiveness (BES-2017/080415 E.C.) and the Spanish Ministry of Science and Innovation (PRE2020/093032 P.M.), and P.T. by a predoctoral fellowship (FPU) from the spanish Ministry of Universities (18/00069). Work was supported by project grants PID2019-110292RB-100 and PID2022-136891NB-I00 (from Spanish Ministry of Science and Innovation), (to J.L.T.).

## Author Contributions

J.L. Trejo designed research; E. Cintado, P. Muela, L. Martín-Rodríguez, I. Alcaide, P. Tezanos, and M. López de Ceballos performed research and analyzed data; K. Vlckova^2,3^, B. Valderrama^2,3^, and T.F.S. Bastiaanssen performed the 16S amplicon microbiota composition analysis, M. Rodríguez-Muñoz performed qPCRs, P. Muela performed the bioinformatic analysis, E. Cintado, P. Muela, M.R. Aburto, J.F. Cryan, María L. de Ceballos, and J.L. Trejo wrote the paper.

## Data availability statement

All materials, data and associated protocols used in this work are available to interested readers. Besides, 16S rRNA sequencing data can be found in European Nucleotide Archive (ENA) under accession PRJEB80752.

## Ethics Approval

All experiments were performed according to the European Community Guidelines (Directive 2010/63/EU) and Spanish Guidelines (Royal Decree 53/2013) and related rules, and they were first validated by the Committee of Ethics and Animal Experimentation of the Cajal Institute (20/05/2016), subsequently favorably evaluated by the CSIC Ethics Committee (Subcommittee of Ethics) of the Spanish Research Council (07/27/2016) and eventually authorized by the competent authority, the Animal Protection Area of the Department of Environment of the Community of Madrid (10/26/2016 and 06/19/2020).

